# Mechanisms of *Pseudomonas aeruginosa* resistance to Type VI Secretion System attacks

**DOI:** 10.1101/2024.10.26.620397

**Authors:** Alejandro Tejada-Arranz, Annika Plack, Minia Antelo-Varela, Andreas Kaczmarczyk, Alexander Klotz, Urs Jenal, Marek Basler

**Author notes:** Current affiliation: Proteomics Unit, Gulbenkian Institute for Molecular Medicine, Lisbon, Portugal (Minia Antelo-Varela). Current affiliation: Department of Biosystems Science and Engineering, ETH Zurich, Basel, Switzerland.

## Abstract

The Type VI Secretion System (T6SS) is a molecular nanomachine that injects toxic effector proteins into the environment or neighbouring cells, playing an important role in interbacterial competition and host antagonism during infection. Pseudomonas aeruginosa encodes three T6SSs. The H1-T6SS delivers toxins in response to attacks mediated by the T6SS of aggressive bacteria, suggesting that P. aeruginosa can resist T6SS assaults. The mechanisms of resistance are poorly characterized. Here, we performed a CRISPRi screen to identify pathways involved in resistance to T6SS effectors of Acinetobacter baylyi and Vibrio cholerae. We show that members of the GacA/GacS regulon, such as the mag operon or aas, and GacA-independent factors, such as the outer membrane protein OprF, confer resistance to different types of T6SS toxins. Interestingly, some of these T6SS resistance mechanisms lead to higher antibiotic susceptibility, suggesting complex evolutionary links between T6SS and antibiotic resistance.

## Introduction

*P. aeruginosa* is a Gram-negative, ubiquitous, opportunistic pathogen that can colonize open wounds and the respiratory airways, and is particularly problematic for patients suffering from cystic fibrosis^1^. *P. aeruginosa* possesses three Type VI Secretion Systems (T6SS), molecular nanoweapons that deliver toxic effectors to neighbouring cells^2^. The first one, called H1-T6SS, plays a predominantly antibacterial role, while the H2- and H3-T6SSs are important for internalization and colonization of a eukaryotic host, although they also deliver trans-kingdom effectors that target both eukaryotic and bacterial cells^2–5^.

Generally, T6SSs are composed of a membrane complex and a baseplate that trigger the assembly of an Hcp tube that is loaded with a tip complex, composed of a VgrG trimer and PAAR proteins, as well as a set of effectors that can be fused to Hcp, VgrG or PAAR or can be loaded separately^6,7^. The Hcp tube is surrounded by a sheath that, upon contraction, propels the Hcp tube as well as the tip complex and all loaded effectors to the extracellular space or into a neighbouring cell. Bacteria are typically protected from their own effectors by a set of dedicated immunity proteins that are encoded adjacently to the effectors^8^. These immunity proteins are highly specific, with few examples of cross-protection to related effectors with similar mechanisms of action^9^.

Interestingly, the H1-T6SS of *P. aeruginosa* was reported to respond to incoming attacks from neighbouring bacteria, regardless of whether they are sister cells or members of a different species^10,11^. Specifically, the H1-T6SS comprises a sensor module, composed by TagQRST, that senses membrane damage and triggers the assembly of a retaliatory H1-T6SS through post-translational modifications^10^. Furthermore, it was proposed that this retaliation strategy is a poor evolutionary strategy, as it prevents *P. aeruginosa* from attacking first and forces it to resist incoming T6SS attacks^12^. Here, we hypothesize that *P. aeruginosa* must be equipped with specific mechanisms that protect against attacks with foreign T6SS effectors, at least until the H1-T6SS is able to strike back.

*Acinetobacter baylyi* ADP1 and *Vibrio cholerae* 2740-80 are Gram-negative environmental bacteria that each encode one antibacterial T6SS cluster with multiple effectors that is highly active *in vitro*. *A. baylyi* ADP1 WT contains four described effectors^13^: Tpe1 (a predicted metallopeptidase without identified antimicrobial activity), Tae1 (a PG-targeting effector), Tse2 (of unknown mechanism of action) and Tle1 (a lipase). *V. cholerae* 2740-80 WT also has four effectors^14^: TseH and VgrG3 (two PG-targeting effectors), TseL (a lipase) and VasX (a pore-forming toxin). Thus, they have different types of effectors, and the resistance mechanisms of *P. aeruginosa* against these two organisms might differ.

In this work, we competed H1-T6SS-deficient *P. aeruginosa* PAO1 carrying a CRISPRi plasmid library targeting all *P. aeruginosa* genes with T6SS-positive or –negative *A. baylyi* ADP1 or *V. cholerae* 2740-80. Deep sequencing of the plasmid library revealed genes required to resist incoming T6SS attacks. Interestingly, most of these genes belong to the regulon of the GacS/GacA two component system, including the previously identified *arc1* and *arc3* clusters and *pa3267*, *aas*^15^, as well as the *mag* operon. The outer membrane protein OprF also contributes to T6SS resistance. Our data suggest that these pathways are general protection mechanisms against multiple effectors and thus differ from specific immunity proteins. In addition, we identified a link between these resistance mechanisms and antibiotic resistance.

## Results

### *P. aeruginosa* resists T6SS attacks in the absence of its H1-T6SS

The H1-T6SS of *P. aeruginosa* can promptly and efficiently retaliate to incoming T6SS attacks, eliminating the competing bacterium before it can efficiently kill *P. aeruginosa* ^10,12,16^. Therefore, we constructed a mutant lacking *tssM1*, a structural membrane component of the H1-T6SS, that is unable to assemble its H1-T6SS and retaliate to incoming attacks. We observed that the H1-T6SS is indeed important to efficiently resist attacks from *A. baylyi* ADP1 (Figure 1A and Supplementary Figure 1A), but not from *V. cholerae* 2740-80 (Figure 1B and Supplementary Figure 1B). Furthermore, although this strain is more susceptible than its parental to attacks from the T6SS of *A. baylyi* ADP1, it is still far more resistant than *E. coli* (Figure 1A and 1C and Supplementary Figures 1A and 1C), as evidenced by the almost complete elimination of *E. coli* when competed in a 10:1 attacker:prey ratio and the strong killing observed in a 1:1 ratio. This suggests that *P. aeruginosa* possesses protective mechanisms against T6SS attacks.

**Figure 1.**
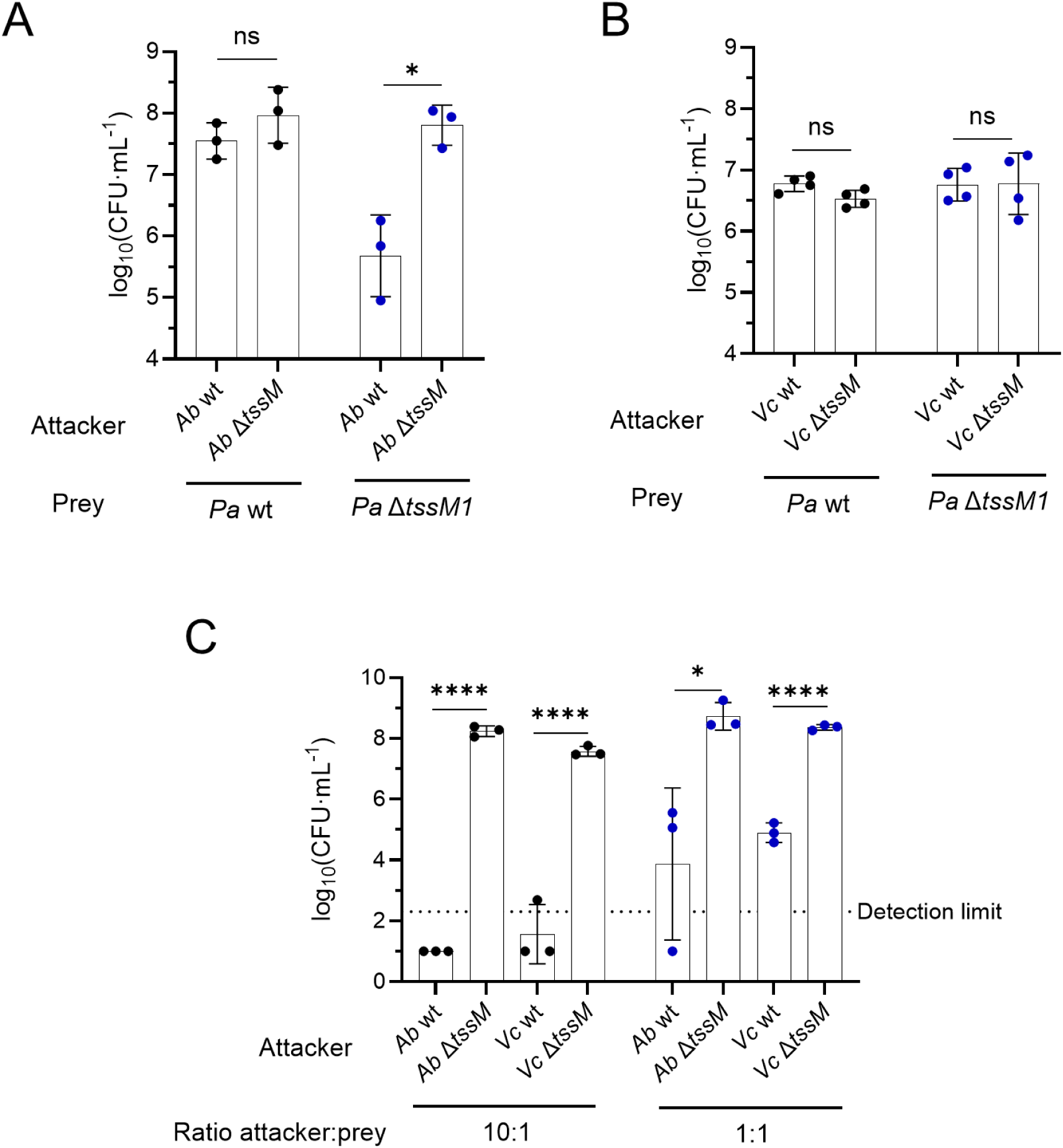
*P. aeruginosa* is more resistant to T6SS from *A. baylyi* and *V. cholerae* than *E. coli*. CFU counts showing the survival of *P. aeruginosa* when competed against *A. baylyi* (*Ab*) (A) and *V. cholerae* (*Vc*) (B) strains that have or not an active T6SS, in a 10:1 attacker:prey ratio where *P. aeruginosa* is the prey. (C) CFU counts showing the survival of *E. coli* (*Ec*) when competed against *A. baylyi* and *V. cholerae* in 10:1 and 1:1 attacker:prey ratios where *E. coli* is the prey.

### Identification of T6SS-protective pathways

In order to identify new T6SS-protective pathways, we introduced the dCas9 protein from *Streptococcus pasteurianus* into *P. aeruginosa* Δ*tssM1* strain and performed a CRISPRi screen for genes important for T6SS protection using a sgRNA library similarly to others^17,18^. Specifically, we competed this T6SS-deficient *P. aeruginosa* strain carrying the sgRNA plasmid library with T6SS-active or inactive mutants (lacking *tssM*) of *A. baylyi* ADP1 or *V. cholerae* 2740-80. We performed two rounds of 3h long incubation of predator and prey at 1:1 and 10:1 ratios, respectively (Figure 2A). The sgRNA loci of the recovered *P. aeruginosa* cells were PCR-amplified (before competition and after the first and second rounds of competition) and the amplicons were sequenced by Illumina sequencing. Samples from competition with T6SS-active or inactive strains were compared in order to identify the pathways that, when knocked down, lead to higher sensitivity to T6SS attacks.

**Figure 2.**
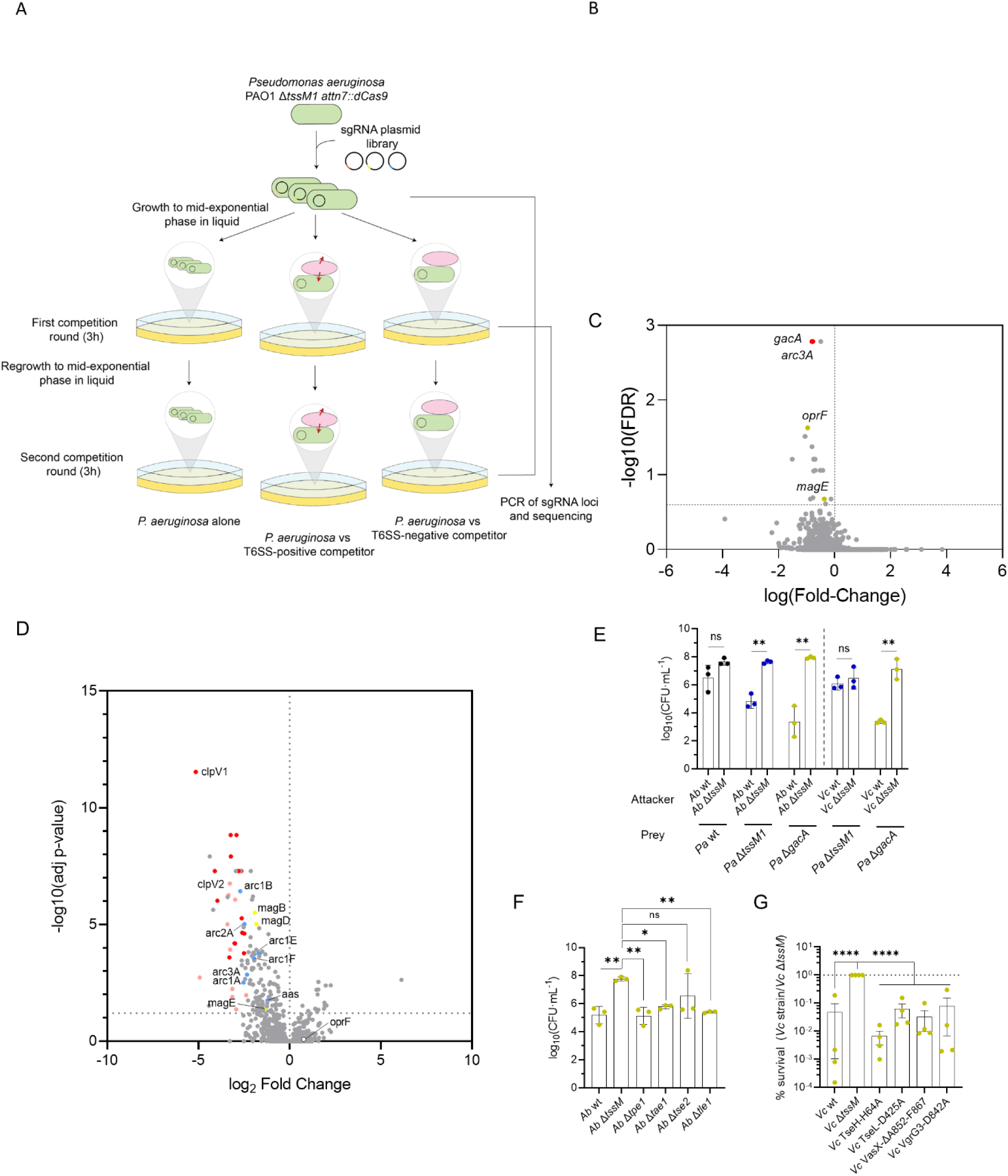
CRISPRi screen for *P. aeruginosa* knockdowns that are sensitive to T6SS attacks. (A) Schematic of the experimental design. *P. aeruginosa* lacking *tssM1*and carrying the sgRNA library was competed against T6SS-positive or –negative *A. baylyi* and *V. cholerae* for 1 or 2 rounds of competition lasting 3h each. Bacteria were grown to mid-exponential phase before each competition round. (B) Negative-selection Volcano plot showing genes important for resistance to T6SS from *V. cholerae*. In red are previously identified genes and in yellow genes newly identified in this study. (C) Negative-selection Volcano plot showing genes important for resistance to T6SS from *A. baylyi*. (D) Volcano plot showing proteins that are differentially regulated in a Δ*gacA* strain compared to wild-type *P. aeruginosa*. In red are components of the H1-T6SS, in pink components of the H2-T6SS, in blue previously identified members of the GacA regulon that are linked to resistance to T6SS attacks of *B. thailandensis*^15^ and in yellow the newly identified hits from the *mag* operon. *oprF* is also highlighted as a non-differentially regulated gene. (E) CFU counts showing the survival of different *P. aeruginosa* strains when competed against T6SS-positive or negative *A. baylyi* and *V. cholerae*. (F) CFU counts showing the survival of *P. aeruginosa* Δ*gacA* when competed against *A. baylyi* lacking different T6SS effectors. (G) Percentage of survival of *P. aeruginosa* Δ*gacA* when competed against *V. cholerae* carrying inactivated versions of different T6SS effectors, compared to *V. cholerae* Δ*tssM*.

After one and two rounds of competition, plasmids encoding sgRNAs targeting a number of *P. aeruginosa* genes were depleted (Figure 2B, C and Supplementary Figure 2A, B). The same sgRNAs were identified in both cases, although differences were more pronounced after two rounds of competition. This included known T6SS resistance-related genes such as *gacA*, *gacS*, *aas*, *arc1A* or *arc3A*^15^ as well as novel genes such as the *mag* operon or *oprF*.

### GacA is a key regulator involved in T6SS resistance

Interestingly, many of the genes that we identified were reported to be under the control of the GacA/GacS two component system (TCS), including *arc1A, arc3A* and *aas*^15^. To determine whether the other genes that we identified are also part of the regulon of the GacA/S TCS, we compared the proteome of a Δ*gacA* strain compared to its parental. We found that the members of the *mag* operon are also downregulated in the Δ*gacA* strain (Figure 2D, Supplementary Figure 2C, and Supplementary Table 1), together with components of the H1- and H2-T6SS, whereas *oprF* is not differentially expressed in that strain.

Next, we assessed whether a Δ*gacA* strain was able to resist T6SS attacks from *A. baylyi* and *V. cholerae*. Indeed, the Δ*gacA* strain was 100-fold more susceptible to T6SS attacks from *A. baylyi* and *V. cholerae* than a strain lacking the H1-T6SS (Figure 2E and Supplementary Figure 2D), indicating that GacA-regulated genes other than the H1-T6SS are required for T6SS resistance.

To get more insights in the mechanisms of resistance, we competed *P. aeruginosa* Δ*gacA* strain with *A. baylyi* strains lacking individual effectors (Figure 2F and Supplementary Figure 2E). We previously showed that these *A. baylyi* strains are able to assemble its T6SS as efficiently as the parental strain^13^. Interestingly, the Δ*gacA* strain is killed similarly when single effectors are absent, although Tse2 might play the most predominant role, as an *A. baylyi* strain lacking *tse2* kills *P. aeruginosa* less efficiently (Figure 2F).

Furthermore, we also competed *P. aeruginosa* Δ*gacA* with *V. cholerae* strains encoding individually inactivated effectors through point mutations that were previously reported^11,19^, and we found that the inactivation of each effector leads to similar levels of killing of the Δ*gacA* strain (Figure 2G and Supplementary Figure 2F). Taken together, our results suggest that the GacA regulon contains genes that contribute to the resistance to different effector classes, in addition to lipase effectors as previously reported^15^.

### The *mag* operon is important for resistance against PG-targeting effectors

Next, we assessed how individual genes from the GacA regulon contribute to T6SS resistance, and in particular the *mag* operon that was identified in our CRISPRi screen (Figure 2B). In *P. aeruginosa*, the *mag* operon is composed of 6 genes, *magABCDEF*. MagD displays weak homology with type II α2-macroglobulin^20^, homologs of eukaryotic α-macroglobulins, whereas the rest of the genes of the operon are important for the folding and stability of MagD^21^. Eukaryotic α2-macroglobulins are innate immunity proteins that protect against proteases by a trapping mechanism^22–24^. However, bacterial type II α2-macroglobulins lack key residues that conform the active site of eukaryotic α2-macroglobulins^21^, and their function is thus unclear.

We constructed a strain lacking both *tssM1* and *magD*, and competed it against different mutants of *V. cholerae*. We performed quantitative killing assays and found that MagD is indeed important for resistance to T6SS attacks, and that it protects mainly against the PG-targeting effector VgrG3 (Figure 3A and Supplementary Figure 3A). To confirm this result, we also performed a CPRG-based lysis assay^13^, a colorimetric assay where lysis is measured based on the conversion rate of the membrane impermeable CPRG dye by a LacZ enzyme that is released upon lysis of *P. aeruginosa*. This assay also indicated a VgrG3-dependent lysis of a Δ*magD* mutant (Figure 3B).

**Figure 3.**
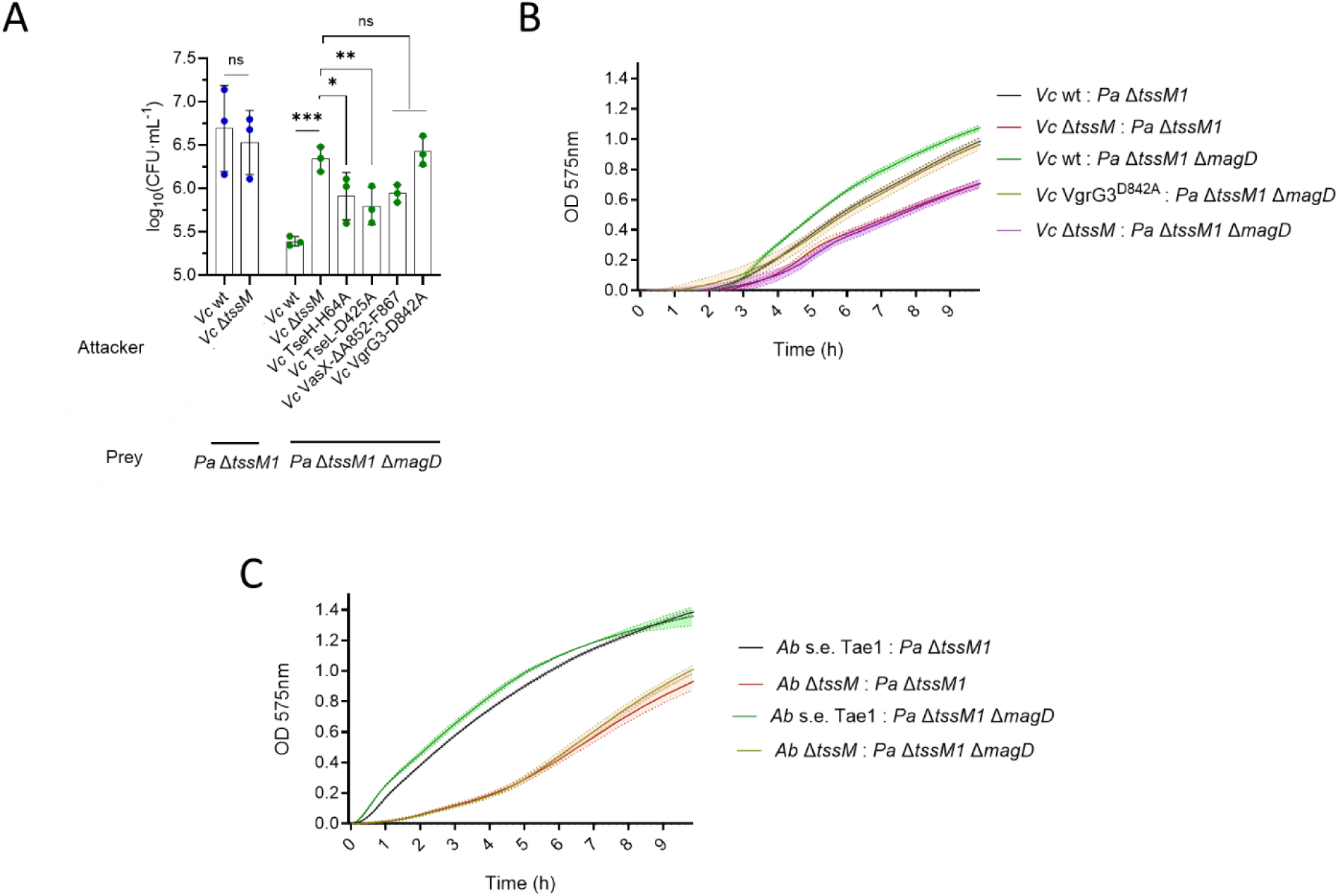
MagD is important for resistance against PG-targeting effectors. (A) CFU counts showing the survival of different *P. aeruginosa* strains when competed against *V. cholerae* carrying different inactivated effectors. (B) CPRG lysis assay showing the lysis levels over time of different *P. aeruginosa* strains when competed against *V. cholerae*. (C) CPRG lysis assay showing the lysis levels over time of different *P. aeruginosa* strains when competed against *A. baylyi* carrying only its PG-targeting effector, Tae1.

Additionally, we tested whether *magD* confers resistance against single effectors of *A. baylyi* by competition with strains expressing single effectors. We found that *magD* also confers a mild protection against the PG-targeting Tae1 effector of *A. baylyi* (Figure 3C), and not against other effector classes (Supplementary Figure 3B-C). Thus, MagD encoded by the *mag* operon specifically protects against PG-targeting effectors from both *V. cholerae* and *A. baylyi*.

### Arc1A, Arc3A and Aas contribute to resistance against lipase effectors

Arc1A and 3A were previously shown to be important for resistance against the T6SS of *B. thailandensis*. Specifically, Arc3A protects against lipase effectors, and the effector class that Arc1A protects against was not clearly elucidated^15^. PA3267 (Aas) was also previously identified in a transposon screen, but no clear role in T6SS resistance was attributed to it, although it was hypothesized that it is important for resistance to lipases^15^. Our screen identified *arc1A*, *arc3A* and *aas*, as important for T6SS resistance. Aas is homologous to the lysophospholipid transporter LplT and the Aas acyltransferase from *E. coli*, where it was shown to be important for defence against phospholipase attacks^25^. We found that a *P. aeruginosa* mutant lacking *tssM1* and either *arc1A*, *arc3A* or *aas* is indeed more sensitive to T6SS attacks from *V. cholerae* WT (Figure 4A). The Δ*tssM1* Δ*arc3A* displays the strongest defect (Figure 4A). A strain lacking all four genes is marginally more sensitive to attacks than the Δ*tssM1* Δ*arc3A* strain, and this sensitivity is abolished when *V. cholerae* lacks an active TseL effector (Figure 4A).

**Figure 4.**
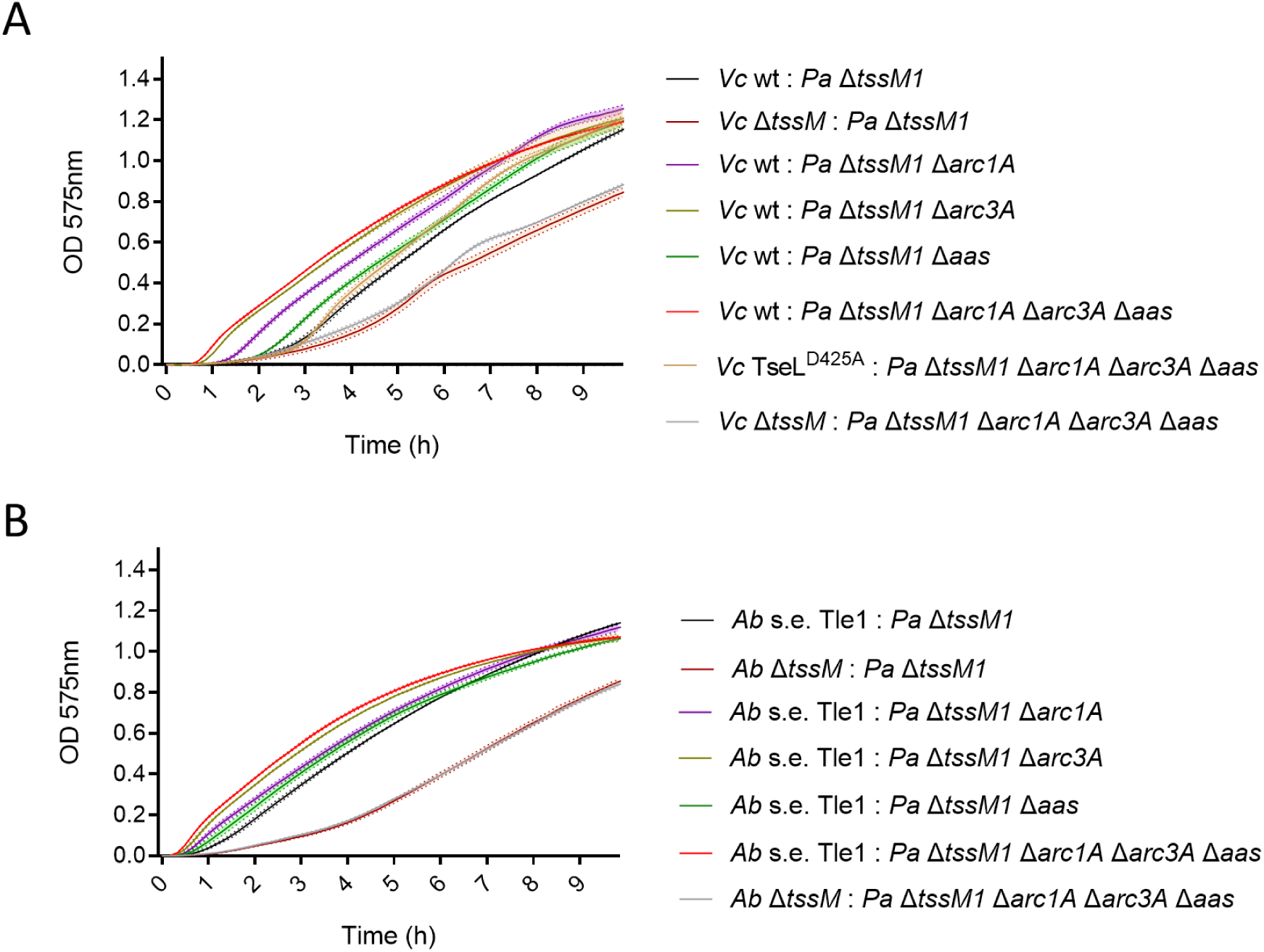
Arc1A, Arc3A and Aas are important for resistance to lipase effectors. CPRG lysis assay showing the lysis levels over time of different *P. aeruginosa* strains when competed against (A) *V. cholerae* and (B) *A. baylyi* carrying only its lipase effector, Tle1.

Similarly, *P. aeruginosa* strains lacking *tssM1* and either *arc1A*, *arc3A* or *aas* are also more sensitive to attacks from *A. baylyi* that carries only its lipase Tle1 effector (Figure 4B). The strain lacking all four genes was not more sensitive to either Tae1 nor Tse2 from *A. baylyi* (Supplementary Figure 3D-E). We conclude *arc1A*, *arc3A* and *aas* are important for resistance to lipase effectors.

### OprF is required for T6SS resistance and is genetically linked to GacA/S

OprF is the main outer membrane porin of *P. aeruginosa*, and it has been reported to perform many different roles including channelling small solutes, maintenance of outer membrane integrity, binding and adhesion, biofilm formation, or outer membrane vesicle biogenesis^26^. OprF is folded in two conformers, a minor population (around 5%) that is folded into a one-domain structure with a channel that acts as porin; and a major population (around 95%) that is folded into two domains including a PG-anchoring domain. It was suggested that this conformation is responsible for anchoring of the OM to the PG^27,28^, and that a Δ*oprF* strain displays defects in the OM^29,30^. To understand how sgRNA-mediated inhibition of *oprF* expression resulted in lower resistance to T6SS attacks, we constructed a *P. aeruginosa* Δ*oprF* strain and competed it against *V. cholerae* and *A. baylyi* attackers. We show that when the attacker strains carry full effector sets, or when single effectors are inactivated, the Δ*oprF* strain is more sensitive than the parental strain (Fig. 5A, B and Supplementary Figure 4A, B). Importantly, Δ*oprF* strain displays a growth defect in LB, which is exacerbated in LB without salt (Supplementary Fig. 4C), suggesting that the strain is sensitive to osmotic shock as previously shown^29^.

**Figure 5.**
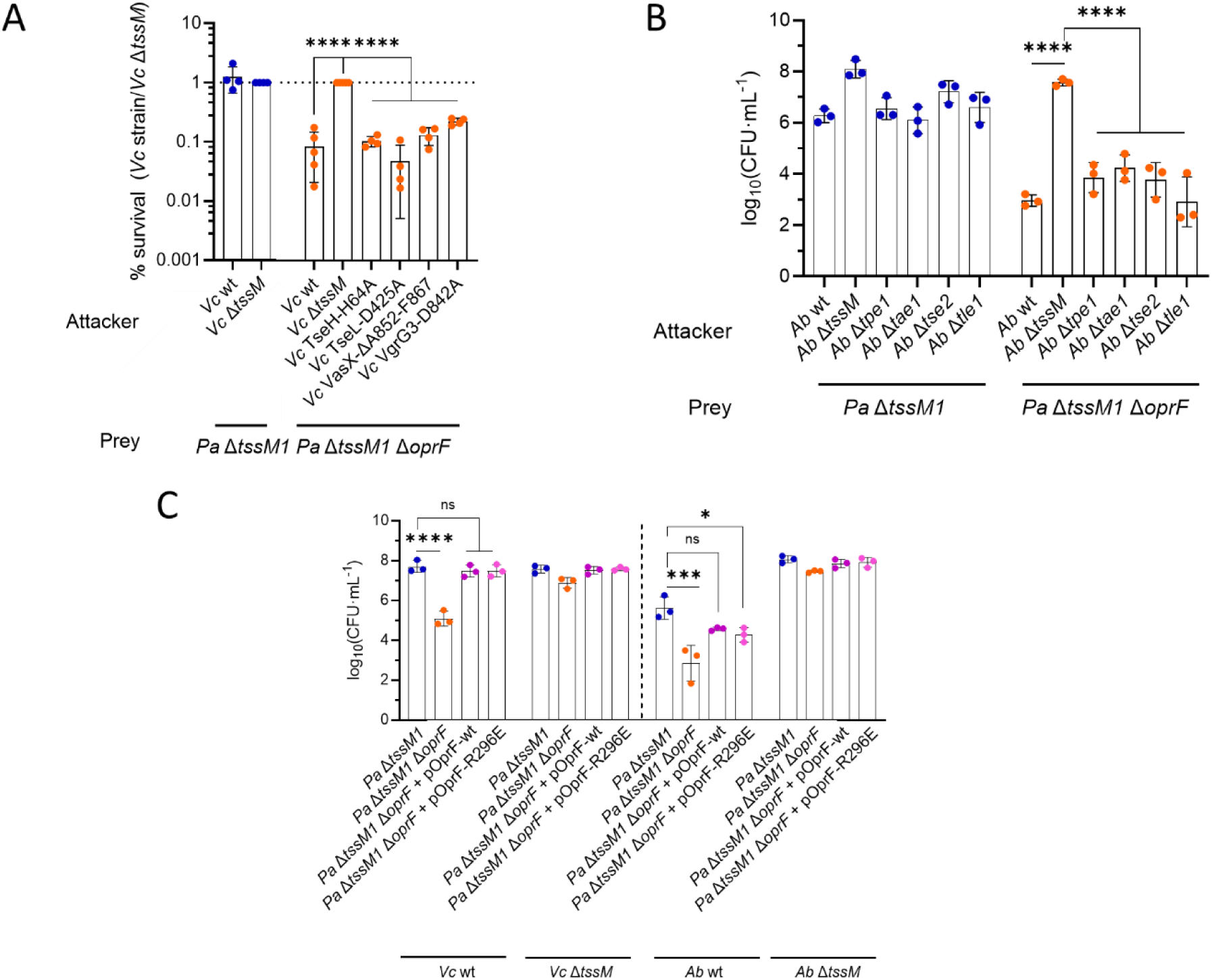
OprF is important for T6SS resistance. (A) Percentage of survival of different *P. aeruginosa* strains with or without *oprF* when competed against *V. cholerae* strains carrying inactivated T6SS effectors, compared to *V. cholerae* Δ*tssM*. (B) CFU counts showing the survival of different *P. aeruginosa* strains when competed against *A. baylyi* strains lacking single T6SS effectors. (C) CFU counts showing the survival of different *P. aeruginosa* strains complemented with wild-type or mutant variants of *oprF* when competed against *V. cholerae* and *A. baylyi*.

Furthermore, complementation *in trans* with a wt copy of *oprF* in a pPSV35 plasmid fully restores both growth and resistance to T6SS attacks (Fig. 5C and Supplementary Fig. 4D and E). The R296E *oprF* variant, that was previously shown to be unable to bind PG in *E. coli*^31,32^, restores resistance to T6SS attacks to the same extent (Fig. 5C and Supplementary Fig. 4E), suggesting that the contribution of the OM anchoring function of OprF to T6SS resistance may be minimal.

Interestingly, we observed different colony sizes when the *P. aeruginosa* Δ*tssM1* Δ*oprF* strain was grown in LB without salt, suggesting the appearance of suppressors (Figure 6A). We isolated four of these clones and analysed their growth, cell morphology and ability to compete with *V. cholerae*. These suppressors displayed wild type-like growth on LB without salt (Figure 6B), as well as wild-type cellular morphology (Figure 6C), as shown by the absence of cell ghosts that are frequent in the Δ*tssM1* Δ*oprF* strain. However, the suppressors were still susceptible to T6SS attacks from *V. cholerae* at a 1:1 ratio (Figure 6D and Supplementary Fig. 4F).

**Figure 6.**
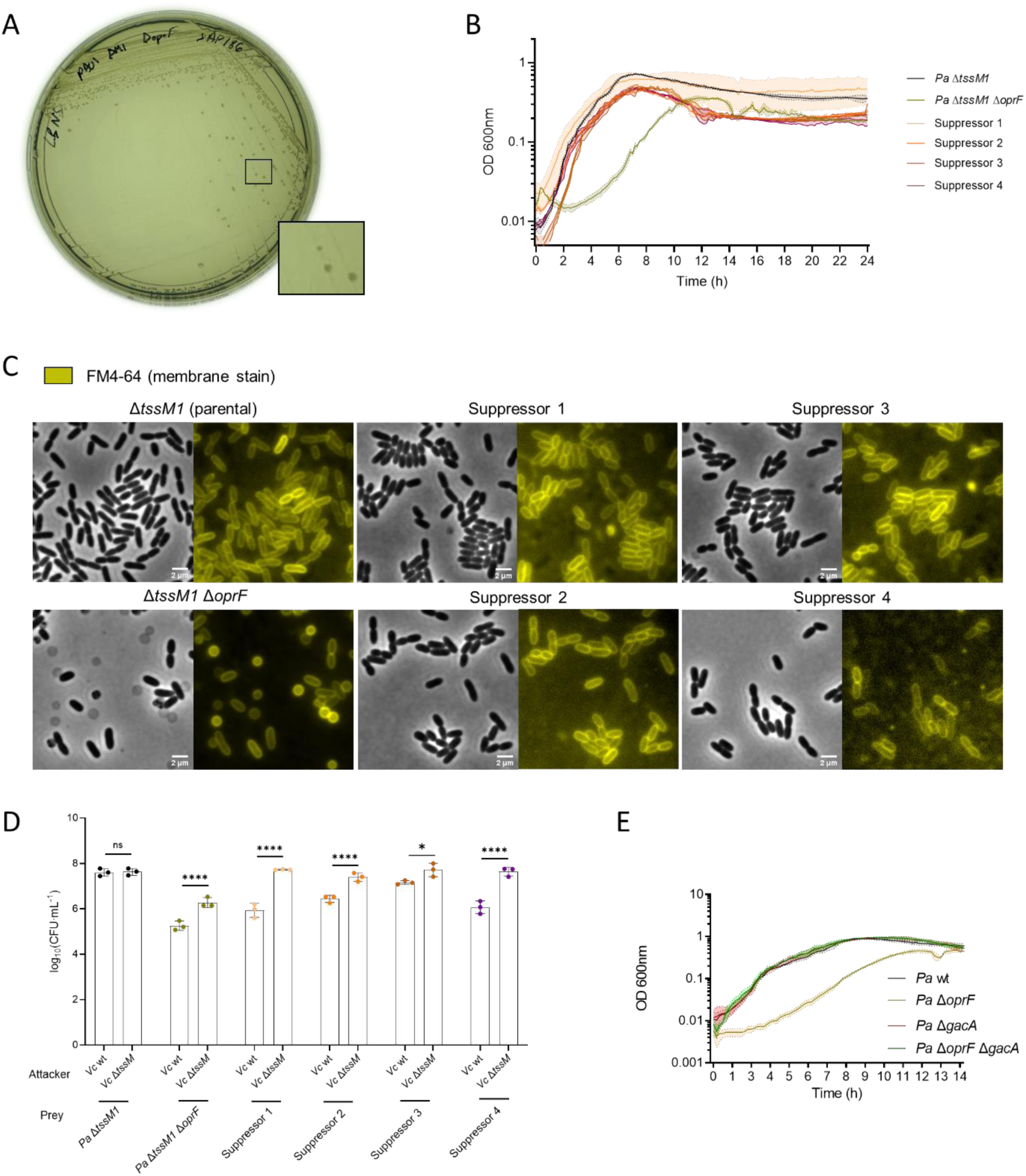
Δ*oprF* suppressors grow normally and recover wild-type morphology but fail to resist T6SS attacks. (A) Image of a *P. aeruginosa* strain lacking *tssM1* and *oprF* growing on an LB no salt plate after overnight incubation at 37°C. (B) Growth curves of parental and suppressor strains of *P. aeruginosa* Δ*tssM1* Δ*oprF*. (C) Fluorescence microscopy images of parental and suppressor strains of *P. aeruginosa* lacking *tssM1* and *oprF*. (D) CFU counts showing the survival of the different *P. aeruginosa* suppressors when competed against T6SS-active or –inactive *V. cholerae*. (E) Growth curve of wild-type and mutant *P. aeruginosa* strains lacking *oprF* and *gacA* or not.

We performed whole genome sequencing of these suppressors and found that all of them carry the same inactivating 431bp deletion within the *gacS* gene, suggesting that the GacA/S is inactive in these strains. We validated this by deletion of *gacA* in a Δ*oprF* strain, which restores the growth defect of this strain (Figure 6E). Thus, GacA/S is involved in causing the growth and morphology defect of the Δ*oprF* strain, which explains why these suppressors are still sensitive to T6SS attacks, as they lack a functional GacA/S that is important for resistance to such attacks (Figure 2). Our results also show that there is a genetic link between *oprF* and the GacA/S TCS.

### Immunity strategies are species-specific

While the previous study from Ting *et al*.^15^ identified three operons, *arc1*, *arc2* and *arc3*, involved in resistance to T6SS from *Burkholderia thailandensis*, only two of them, *arc1* and *arc3* appeared in our CRISPRi screen. Likewise, the two novel resistance factors that we identified, the *mag* operon and *oprF*, were not identified in the previous transposon screen. In order to verify whether these mechanisms are important for resistance to other organisms, we also tested the resistance of our strains to *B. thailandensis* attacks (Fig. 7 and Supplementary Figure 5). Although *magD* is dispensable for resistance to the T6SS of *B. thailandensis*, both *gacA* and *oprF* play a role, albeit to a lesser extent than in the case of *A. baylyi* or *V. cholerae*. Thus, some T6SS-protective mechanisms, such as those in the GacA regulon, are specific to certain species or effector types, and others, such as OprF-mediated mechanisms, appear to be generic resistance mechanisms.

**Figure 7.**
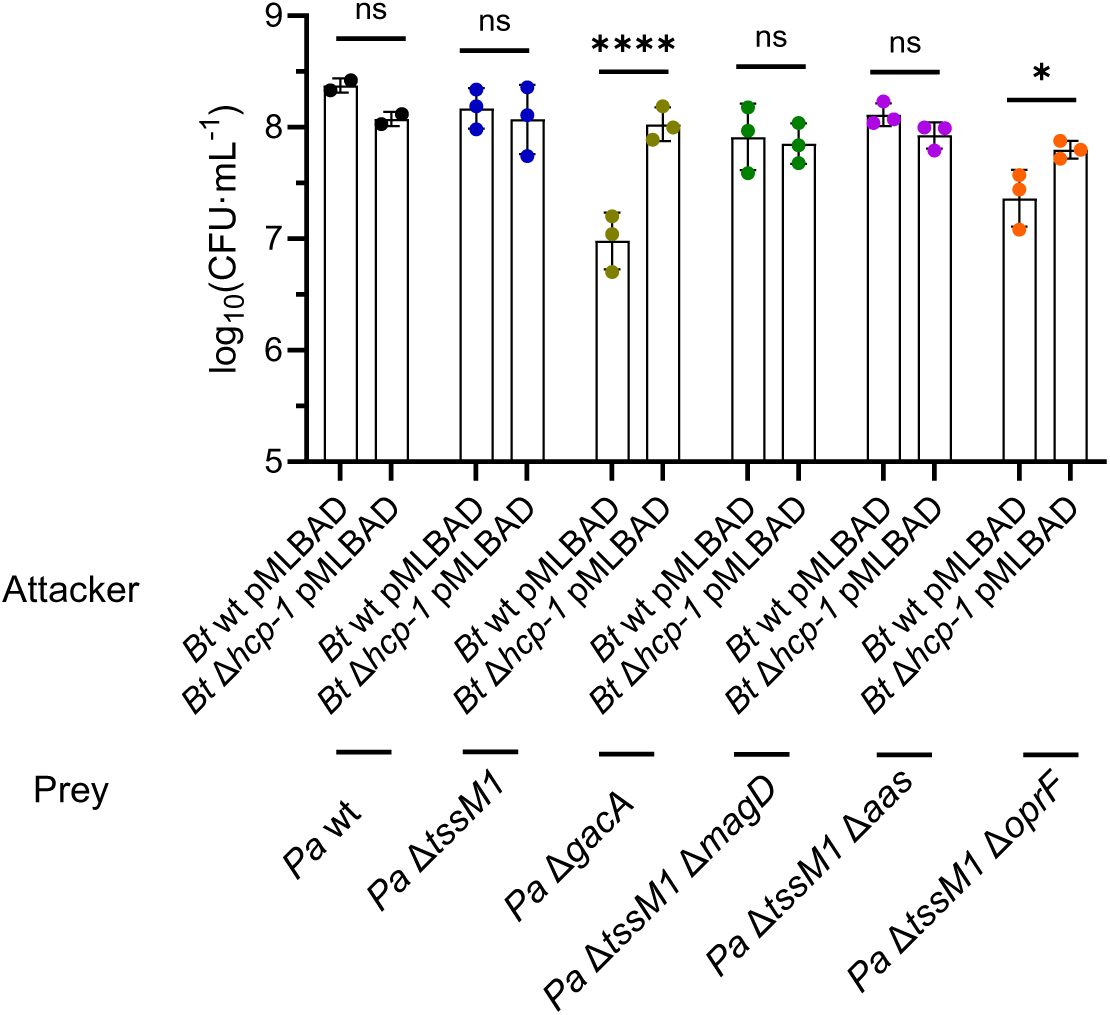
T6SS resistance mechanisms are species- or effector-specific. CFU counts showing the survival of *P. aeruginosa* strains lacking different T6SS resistance mechanisms when competed with *B. thailandensis* (*Bt*) in a 10:1 attacker:prey ratio where *P. aeruginosa* is the prey.

### T6SS immunity strategies can affect antibiotic resistance

To determine whether T6SS resistance mechanisms might be important for antibiotic resistance, we explored the susceptibility of T6SS-sensitive strains to 30 antibiotics from different classes, including inhibitors of translation, cell wall synthesis, folate synthesis, membrane integrity, DNA replication and transcription, among others. We performed growth curves of the Δ*magD*, the Δ*tssM1* Δ*arc1A* Δ*arc3A* Δ*aas* and the Δ*oprF* strains in the presence of antibiotics and compared the cumulative OD_600 nm_ between said conditions, relative to WT and to the untreated condition (Figure 8A and Supplementary Figure 6). Interestingly, all three tested strains displayed altered antibiotic susceptibility profiles compared to WT. *P. aeruginosa* Δ*magD* is more resistant to chlorhexidine, an antimicrobial targeting the bacterial cell membrane (Figure 8A-B). The *P. aeruginosa* Δ*tssM1* Δ*arc1A* Δ*arc3A* Δ*aas* strain grows generally better in the presence of fluoroquinolones, such as nadifloxacin or ciprofloxacin (Figure 8A, 8C-D), and merbromin. Lastly, the *P. aeruginosa* Δ*oprF* strain is more resistant to antibiotics like the membrane-targeting colistin (Figure 8A, 8E), and more sensitive to the translation inhibitor tetracycline (Figure 8A, 8F) and beta-lactams such as piperacillin or azlocillin (Figure 8A). All three strains displayed a mild advantage in the presence of gentamicin, that was particularly pronounced in the Δ*oprF* strain (Figure 8A). This suggests that T6SS immunity mechanisms alter the antibiotic susceptibility profile of *P. aeruginosa*.

**Figure 8.**
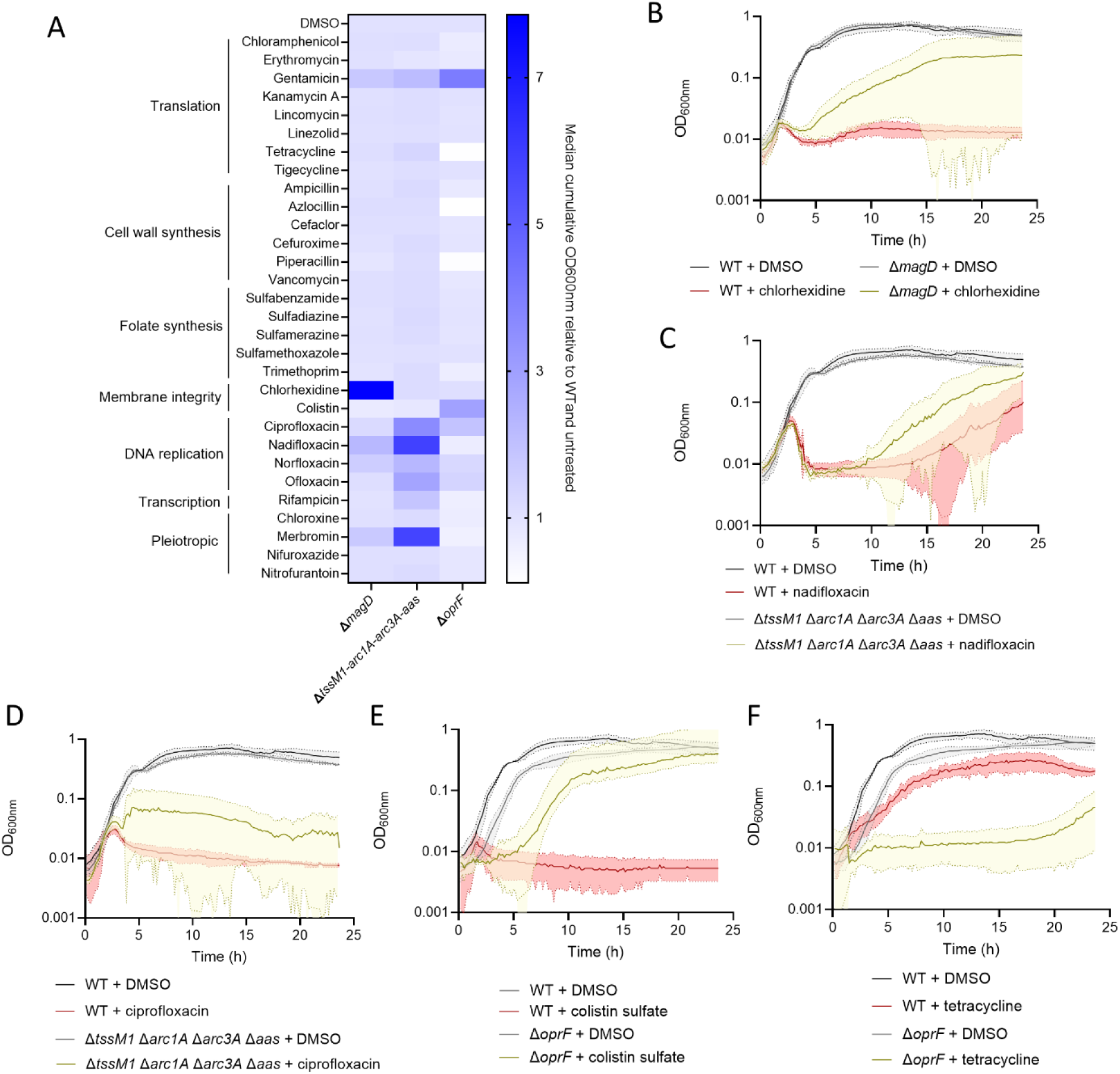
T6SS resistance mechanisms influence antibiotic resistance. (A) Heat-map showing the median cumulative OD_600 nm_ after 23.5h of growth at 37°C, relative to the WT and the untreated condition (DMSO control). (B) Growth curve of *P. aeruginosa* wild-type or Δ*magD* with or without 20 μM chlorhexidine. (C) Growth curve of *P. aeruginosa* wild-type or Δ*tssM1* Δ*arc1A* Δ*arc3A* Δ*aas* with or without 20 μM nadifloxacin or (D) ciprofloxacin. (E) Growth curve of *P. aeruginosa* wild-type or Δ*oprF* with or without 20 μM colistin sulfate or (F) tetracycline.

## Discussion

*P. aeruginosa* is able to promptly sense and retaliate to T6SS attacks from other species^10^. However, the mechanisms that allow *P. aeruginosa* to withstand the initial attack are not well understood. Only one recent report^15^ identified three operons involved in resistance to T6SS attacks from *B. thailandensis*, called *arc1*, *arc2* and *arc3*, and determined that *arc3* is particularly important to resist the delivery of lipase effectors, likely through the detoxification of lysophospholipids that are generated as a result of the activity of these lipases. However, the effector types that *arc1* and *arc2* confer resistance against, and whether there are other mechanisms involved in the resistance to other effector types, is not known. Here, we performed a CRISPRi screen to identify novel factors involved in T6SS resistance in *P. aeruginosa*, using the laboratory models *A. baylyi* and *V. cholerae* as aggressor species. These two species contain different effector sets^13,14^: both of them have at least one lipase (Tle1 in *A. baylyi* and TseL in *V. cholerae*) and at least one PG-targeting effector (Tae1 in *A. baylyi* and VgrG3 and TseH in *V. cholerae*); *A. baylyi* encodes two other effectors of unknown activity (Tpe1 and Tse2); and *V. cholerae* encodes one membrane-targeting pore-forming effector VasX.

In our screen, we identified known factors such as the *arc1* and *arc3* clusters, as well as *gacS* and *gacA* of the GacA/S TCS that regulates the expression of said clusters^15^ in addition to the expression of the H1- and H2-T6SS^33,34^ (Figure 2B-C). We also identified novel factors that contribute to resistance, the *mag* operon, that is also regulated by GacA, underscoring the importance of this transcriptional regulator for the survival of *P. aeruginosa* in polymicrobial communities. Furthermore, we found other factors that are not known to be regulated by GacA (Figure 2E), such as the outer membrane protein OprF.

Through quantitative killing assays and kinetic CPRG-based lysis assays, we could validate that these factors are indeed involved in resistance to T6SS attacks (Figure 2-5). In agreement with Ting *et al.*^15^, we also observed that *arc3* is important specifically to resist to lipases (Figure 4). Here, we also observe that *arc1* as well as *aas* are required for full resistance against the lipases from both *V. cholerae* and *A. baylyi*. In their study, Ting *et al*. did not determine the effector type that *arc1* is active against and, although they hypothesized that *aas* is active against lipases^15^, they did not find a clear phenotype against *B. thailandensis.* This suggests that the specific catalytic mechanism of the lipases in question might be relevant for the mechanism of action of the corresponding resistance mechanisms.

Our results also show that the GacA regulon is important for the resistance against effector classes other than lipases. We found that the *mag* operon, that is regulated by GacA, is particularly important to resist against certain PG-targeting effectors such as VgrG3 from *V. cholerae* and Tae1 from *A. baylyi*. Interestingly, the central gene of this operon, *magD*, displays homology with human α2-macroglobulins^20^, innate immunity factors that defend against proteases through a trapping mechanism^22–24^. However, the catalytic residues of this protein family are not conserved in bacterial type II α2-macroglobulins that *magD* belongs to^21^, raising questions about the true function of this protein family in bacteria. Here, we found that MagD contributes to resistance to T6SS PG-targeting effectors, although the exact molecular mechanism remains to be elucidated. Interestingly, MagD was previously found to co-immunoprecipitate with TssM1^20^, a structural transmembrane component of the H1-T6SS of *P. aeruginosa*, suggesting that these two components might co-localize where a foreign T6SS attack was detected and contribute to the defense against the injected effectors (in the case of MagD) and the assembly of a counterattack H1-T6SS machine (in the case of TssM1).

*P. aeruginosa* also possesses other resistance mechanisms that are not known to be regulated by GacA (Figure 2D), such as the outer membrane protein (OMP) OprF. OprF is the most abundant OMP in *P. aeruginosa*, and carries out multiple functions including an outer membrane porin as well as anchoring the outer membrane to the PG^26^. Here, we found that OprF contributes to resistance to multiple T6SS effector classes (Figure 5A and B and Supplementary Figures 4A and B). We confirmed this phenotype by complementation with a wild-type or mutant variant of OprF that no longer binds PG (Figure 5C), and found that this variant is similarly able to promote survival against both *V. cholerae* and *A. baylyi*, suggesting that other functions of OprF besides OM anchoring must be important for survival to T6SS attacks.

Our study also provides the first evidence for the genetic relationship between *oprF* and the *gacA/S* TCS. A strain lacking *oprF* displays a growth defect in LB that is more pronounced in LB without salt (Supplementary Figure 4C). In LB without salt, this strain spontaneously reverts to wild-type-like growth at high frequency (Figure 6A and B) and, although these colonies also display WT-like morphology (Figure 6C), they are still unable to survive T6SS attacks from *V. cholerae* (Figure 6D). Our whole genome sequencing analysis showed that these suppressors carry a loss-of-function deletion of *gacS*, which explains why these strains are susceptible to T6SS attacks. Indeed, the GacA/S TCS is responsible for the growth defect of the Δ*oprF* strain, since deletion of *gacA* restores this defect (Figure 6E). Further research is needed in order to determine why the expression of the GacA regulon leads to a growth defect in the Δ*oprF* genetic background.

Next, we also assessed whether our newly identified resistance mechanisms also play a role against *B. thailandensis* E264. As expected from a transposon screen performed by Ting *et al*.^15^, *magD* does not play a role in defending from attacks from *B. thailandensis*. However, both GacA and OprF are important for this (Figure 7 and Supplementary Figure 5). Interestingly, the *arc2* operon identified by Ting *et al*.^15^ did not appear in our screen. Taken together, our results show that *P. aeruginosa* possesses an array of T6SS resistance mechanisms, frequently under the control of the GacS/A TCS, that help it survive attacks from a variety of aggressor species that it can encounter in the environment. However, these resistance mechanisms might only protect against certain species or against effectors with specific modes of action.

Interestingly, other general stress response factors such as those previously found to protect *E. coli* and *P. aeruginosa*^35,36^ were not highlighted in our screen. In the case of *P. aeruginosa*, those factors were identified upon overexpression of TseL in the host periplasm, a condition that is likely more deleterious for *P. aeruginosa* than a single T6SS attack where only a few molecules of TseL may be injected at the same time^35^. This would suggest that members of the GacA regulon mount a first layer of defence and that, when they are overwhelmed, general stress response mechanisms of the cell come into play. Performing a similar screen in a Δ*gacA* genetic background where dedicated protection mechanisms are not expressed might highlight such general stress response mechanisms.

We also wondered whether these T6SS resistance mechanisms could be involved in resistance to other antimicrobials such as antibiotics. We screened a panel of 30 antibiotics against the WT, Δ*magD*, Δ*tssM1* Δ*arc1A* Δ*arc3A* Δ*aas* and Δ*oprF* and strains (Figure 8 and Supplementary Figure 6) and found that, surprisingly, strains lacking these T6SS resistance mechanisms display a slight growth advantage in the presence of selected antibiotics as compared to WT (chlorhexidine for Δ*magD*; fluoroquinolones and merbromin for Δ*tssM1* Δ*arc1A* Δ*arc3A* Δ*aas*; colistin for Δ*oprF*; and gentamicin for all three strains), suggesting that these T6SS resistance mechanisms may worsen the toxicity of selected antibiotics. As a consequence, we hypothesize there are trade-offs to obtaining T6SS resistance mechanisms that may lead to higher antibiotic susceptibility in *P. aeruginosa*. Similar trade-offs were also identified in *E. coli* that evolved resistance against T6SS attacks from *V. cholerae*^37^.

It is known that colistin kills Gram-negative bacteria by targeting LPS and disrupting the OM^38,39^. Most mechanisms of resistance to colistin have been linked to LPS modification^38^. More research is needed to clarify why an *oprF* mutant, that displays defects in OM stability, is more resistant to colistin. On the other hand, a Δ*oprF* strain is also more susceptible to antibiotics such as the ribosome-targeting tetracycline, or the beta-lactams piperacillin or azlocillin. These phenotypes cannot be explained by the porin function of *oprF*^26^, since the strain should be more resistant to these antibiotics if *oprF* was required for drug uptake. Overall, our findings serve as proof-of-concept that the study of the mechanisms of resistance to the T6SS can lead to insights about antimicrobial resistance in bacteria.

In conclusion, in this work we identified multiple novel resistance mechanisms in *P. aeruginosa* against T6SS attacks from *A. baylyi* and *V. cholerae*, which are also linked with resistance to classical antimicrobials. Interestingly, some T6SS resistance mechanisms may lead to increased sensitivity to antibiotics, suggesting that these resistance mechanisms may not be always beneficial. In the future, further research will be needed to determine whether these resistance mechanisms play a role in the survival of *P. aeruginosa* strains in naturally-occurring polymicrobial communities, and particularly in the context of clinical infections, and in the presence of antibiotics.

## Materials and methods

### Bacterial strains and growth conditions

Bacterial strains used in this study can be found in Supplementary Table S2. Bacteria were grown in LB at 37°C unless otherwise stated. *E. coli* DH5α λpir was used as cloning strain and *E. coli* SM10 λpir was used for conjugation of *P. aeruginosa* and *V. cholerae*. Antibiotics used were streptomycin (100 μg/mL for *V. cholerae* and *A. baylyi* on solid media; and 50 μg/mL for *A. baylyi* in liquid media), gentamicin (20 μg/mL for *P. aeruginosa* and *E. coli*), kanamycin (50 μg/mL for *A. baylyi*), carbenicillin (150 μg/mL for *E. coli* and *V. cholerae* and 100 μg/mL for *P. aeruginosa*), irgasan (20 μg/mL for *P. aeruginosa*) and trimethoprim (200 μg/mL for *B. thailandensis* and 50 μg/mL for *E. coli*).

### Mutant construction

All plasmids used in this study are listed in Supplementary Table S3, and primers in Supplementary Table S4. *P. aeruginosa* mutants were constructed as previously described using the pEXG2 suicide plasmid^40^. Briefly, 800bp homology arms were amplified by PCR using primers (a list of all primers can be found in Supplementary Table S3) with 20bp stretches that were homologous to the *Hind*III and *Xba*I restriction sites of the pEXG2 plasmid. PCR products were inserted into linearized pEGX2 plasmid by Gibson assembly^41^ (NEB, E2611L). The resulting vectors were transformed into chemically-competent *E. coli* DH5α λpir, selected on LB plates with 20 μg/mL gentamicin and verified by colony PCR and Sanger sequencing. The purified plasmids were then transformed into chemically-competent *E. coli* SM10 λpir and the resulting strains were used for conjugation of *P. aeruginosa* PAO1. Selection of *P. aeruginosa* transconjugants was performed on LB with 20 μg/mL gentamicin and 20 μg/mL irgasan. The pEXG2 backbone was then curated by restreaking on LB plates without salt supplemented with 20 μg/mL irgasan and 10% sucrose, which were incubated for 36h at room temperature. Colonies were screened by colony PCR with flanking primers.

pPSV35-derived plasmids were introduced into *P. aeruginosa* by electroporation of 20-200ng of plasmid into 1 mL of overnight stationary phase cultures washed twice with sterile water using a Bio-Rad GenePulser with the following settings: 25 μF, 400 Ω, 1.8 kV. All steps were performed at room temperature. 1 mL of LB medium was added immediately after the pulse and cells were incubated 1.5-2h at 37°C with shaking before plating on LB plates containing 20 μg/mL gentamicin.

The chromosomal insertion of *dCas9* in the attnT7 site was performed as previously described (Kaczmarczyk *et al.*). Briefly, an 18-22 h old culture of *P. aeruginosa* Δ*tssM1* in LB without salt was washed twice with sterile water and resuspended in 100 μL sterile water. The bacterial suspension was electroporated with two plasmids at the same time, pUC18T:TNT7:dCas9Spas and pTNS2 that contains the Tn7 integrase and is unable to replicate in *P. aeruginosa*. The electroporation was performed as described above. The cells were recovered for 1h in LB at 37°C and 100 μL were plated on LB agar plates with 20 μg/mL gentamicin to select for transconjugants that integrated *dCas9* into the chromosome. The next day, colonies were grown in LB no salt and, as above, electroporated with plasmid pFLP2 containing the flippase that removes the backbone of the pUC18T plasmid. After recovery, 100 μL were plated on LB agar plates with 100 μg/mL carbenicillin and incubated overnight at 37°C. The resulting colonies were restreaked on LB no salt + 10% sucrose to counterselect pFLP2. The successful insertion of *dCas9* was verified by PCR and the resulting strain was phenotypically validated by expression of an *ftsZ*-targeting sgRNA that led to cell filamentation as described^18^.

*V. cholerae* mutants were constructed in a similar manner, using the pWM91 suicide plasmid and the XbaI and HindIII restriction sites. *V. cholerae* transconjugants were selected on LB plates with 150 μg/mL carbenicillin and 100 μg/mL streptomycin. The pWM91 backbone was curated by restreaking on LB plates without salt supplemented with 100 μg/mL streptomycin and 10% sucrose.

*A. baylyi* mutants were constructed by natural transformation as previously described^13^, using a kanamycin resistance, streptomycin counterselectable, cassette and 800bp homology arms.

*B. thailandensis* were conjugated with pMLBAD plasmid as previously described^42^.

### Quantitative killing assays

Overnight cultures of *P. aeruginosa*, *V. cholerae*, *E. coli* and *B. thailandensis* were grown in LB at 37°C and 200 rpm without antibiotics unless *P. aeruginosa* or *B. thailandensis* contained plasmids. *A. baylyi* overnight cultures were grown at 30°C in similar conditions without antibiotics. Day cultures were launched in LB at the following dilutions: 1:100 for *P. aeruginosa*, 1:50 for *A. baylyi* and *B. thailandensis*, 1:200 for *E. coli* and 1:400 for *V: cholerae*. After 2.5-3h, when all cultures reached an OD_600nm_ of 0.7-1.4, cultures were spun down at 12,000g for 2 min at room temperature. The cultures were concentrated to OD 10 with LB and 1:10 prey:attacker mixes were prepared. Five μL of the resulting mixes were spotted on LB agar plates, in technical triplicates, and allowed to dry. The plates were incubated at 37°C for 3-4h and then the spots were excised and resuspended in 500 μL liquid LB by vortexing. Ten-fold serial dilutions were performed and 5 μL of the resulting suspensions were spotted on selective plates. As selective agents, we used irgasan 20 μg/mL for *P. aeruginosa*, streptomycin 100 μg/mL for *V. cholerae* and *A. baylyi*, gentamicin 20 μg/mL for *E. coli* and 200 μg/mL trimethoprim for *B. thailandensis*. Plates were incubated overnight at 37°C and the surviving CFUs were counted the next day and technical triplicates were averaged. The experiments were performed with at least three biological replicates and plotted with Prism.

### CRISPRi library electroporation

A *P. aeruginosa* Δ*tssM1* strain containing *dCas9* at the attnT7 site was grown overnight in LB without salt and 4 mL were electroporated with 2 μL of the CRISPRi library as described above. A manuscript including details on the library composition and experimental validation is currently under preparation (Kaczmarczyk *et al.*). After electroporation, the bacterial suspension was resuspended in 1 mL LB and transferred to a 1 L Erlenmeyer flask containing 100 mL LB and allowed to recover for 1-1.5h at 37°C while shaking. Next, 20 μg/mL gentimicin were added and the culture was incubated overnight at 37°C while shaking. This protocol led to an electroporation efficiency of 3.8·10^6^ cells/mL, which allows to achieve a good coverage of the ca.83,000 sgRNA library.

### CRISPRi screening competition

Three 100 mL day cultures in LB+20 μg/mL gentamicin+1mM IPTG were launched by inoculation of 1 mL at OD_600nm_ 1 from the library electroporation culture. In parallel, 20 mL day cultures were launched from overnight cultures of *V. cholerae* and *A. baylyi* strains. The cultures were incubated for around 3h until an OD_600nm_ of 0.7-1.4 was reached. Five mL of each *P. aeruginosa* culture were taken, pelleted and frozen at −80°C until further processing (time 0 before the competition started). For each replicate, the cultures were concentrated to OD 10 and mixed in a 10:1 *V. cholerae*:*P. aeruginosa* ratio or in a 1:1 *A.baylyi*:*P. aeruginosa* ratio and 90 μL of the resulting mixes were spotted on LB+1mM IPTG plates. *P. aeruginosa* alone was also spotted (no co-incubation control). The spots were allowed to dry and were then incubated at 37°C for 3h. Then, the spots were excised and resuspended in 7 mL LB by vortexing and pipetting. The resuspended bacteria was spun down for 10 min at 3,900g and resuspended in 7 mL LB. Five mL were pelleted and frozen at −80°C until further processing (samples after the first round of competition). The remaining 2 mL were used to measure the OD_600nm_ and inoculate new day cultures in 100 mL LB + 20 μg/mL gentamicin + 1 mM IPTG with 1 mL OD 1. New day cultures of *V. cholerae* and *A. baylyi* were launched similarly as earlier. The same process was repeated for a second round of competition and the resulting samples were again pelleted and frozen at −80°C until further processing.

### CRISPRi sequencing library preparation

Pellets were thawed and resuspended in 500 μL 50 mM NaOH. The samples were incubated for 10 min at 50°C and 1 μL was used as template for a PCR reaction using Phusion DNA Polymerase (M0530S, NEB) as described (Kaczmarczyk *et al.*) using primer pairs oATA402+403, oATA404+405, oATA406+407 and oATA408+409 for a fourth of the samples each. The PCR cycles were performed as follows: 98°C for 3min, 35 cycles of 98°C denaturation for 10 s, 63°C annealing for 10 s and 72°C elongation for 10 s, followed by a final extension at 72°C for 1 min and hold at 10°C until further processing. The PCR products were run on a 2% agarose gel and purified from gel using a NucleoSpin Gel and PCR Clean-up kit (Macherey-Nagel). The DNA concentration of the samples was measured and samples were diluted to 1 ng/μL in 10 μL, of which 5 μL were used as template for the indexing PCR.

Indexing was performed with the kit IDT for Illumina Nextera DNA UD Index Set C (ref 20026934) with a PCR mix as follows: 5 μL DNA template, 5 μL primer mix, 5 μL Phusion GC Buffer, 0.5 μL 10mM dNTPs, 0.25 μL Phusion DNA polymerase (M0530S, NEB) and 9.25 μL H_2_O. The following program was used: 98°C for 3 min, followed by 8 cycles of 98°C denaturation for 10 s, 55°C annealing for 10 s and 72°C elongation for 10 s, and then a final extension at 72°C for 1 min and hold at 10°C until further processing. The PCR products were purified using 1.8 volumes of NEBNext Sample Purification Beads (E7767S, NEB). Samples were resuspended in 20 μL H_2_O and a quality check was run on a TapeStation 4150 device (Agilent) using DNA ScreenTape chips (Agilent) as recommended by the manufacturer. Sequencing was performed as previously described (Kaczmarczyk *et al.*) by Illumina sequencing at the Genomics Facility Basel (ETH Zürich, University of Basel) using a NextSeq SR 81/10 kit.

Data analysis was performed using the European Galaxy server^43^ using the previously described MAGeCK pipeline^44^ using median normalization, FDR-adjusted threshold of 0.25, FDR as p-value adjustment method and 10 as remove zero threshold. The median method as used for the gene log-fold change. Data was plotted using Prism.

### Sample preparation for proteomics

Overnight cultures of *P. aeruginosa* were grown at 37°C and 200 rpm. Day cultures in LB were launched (1:100 dilution of the overnight) and incubated for 2.5-3h at 37°C and 200 rpm, until they reached an OD_600nm_ of 0.7-1.4. They were concentrated to an OD of 10 and 20 μL were spotted on LB plates. The spots were allowed to dry and were then incubated at 37°C for 3 h. The spots were then resuspended in 500 μL phosphate-buffered saline (PBS), pelleted by centrifugation at 12,000g for 2 min and stored at −80°C until further processing. Cell pellets were resuspended in lysis buffer (5% Sodium dodecyl sulfate (SDS), 10mM tris(2-carboxyethyl) phosphine (TCEP), 100 mM Triethyloammonium bicarbonate (TEAB)) followed by incubation at 95°C for 10 min. Cells were disrupted by ultrasonication using the PIXUL system (Active Motif) for 20 min with default settings (pulse 50 cycles, PRF 1 kHz, burst rate 20 Hz)) and the protein content was determined by tryptophan-based fluorescence assay (Infinite M Plex, Tecan). Sample alkylation was performed by addition of 20 mM iodoacetamide (IAA) and incubation at 25°C for 30 min with gentle shaking. 10 μg of protein lysate was digested (trypsin 1/100, w/w; Promega) and purified using the SP3 protocol^45^. Peptide concentration was determined using an UV-based assay (Infinite M Nano, Tecan). Samples were then resuspended at a final concentration of 250 ng/μL.

### SWATH-MS data acquisition

Samples were separated on a Dionex UltiMate 3000 system (ThermoFisher Scientific) coupled online to an Orbitrap Exploris 480 mass spectrometer (ThermoFisher Scientific). In-house packed 20 cm, 75 μm ID capillary column with 1.9 μm Reprosil-Pur C18 beads (Dr. Maisch, Ammerbuch, Germany) was used. The column temperature was maintained at 60 °C using an integrated column oven interfaced online with the mass spectrometer. Formic acid (FA) 0.1% was used to buffer the pH in the two running buffers used. The total gradient time was 60 min and went from 2% to 12% acetonitrile (ACN) in 5 min, followed by 45 min to 35%, and 10 min at 50%. This was followed by a washout by 95% ACN, which was kept for 20 min, followed by re-equilibration of 0.1% FA buffer. Flow rate was kept at 300 nL/min. Spray voltage was set to 2500 V, funnel RF level at 40, and heated capillary at 275 °C. For DIA experiments full MS resolutions were set to 120,000 and full MS normalized AGC target was 300% with an IT of 45 ms. Mass range was set to 350–1400. Normalized AGC target value for fragment spectra was set at 1000%. In all, 63 windows of 9 Da were used with an overlap of 1 Da. Resolution was set to 15,000 and IT to 22 ms. Normalized CE was set at 28%. All data were acquired in centroid mode using positive polarity and advanced peak determination was set to on.

### SWATH-MS data analysis

For data processing and protein identification, raw data were imported into SpectroNaut (16.1.220730.53000, Biognosys) and analyzed with directDIA. Searches were carried out against a fasta file including the proteomes of *P. aeruginosa* (UP000002438). Cysteine carbamidomethylation was set as fixed modification. Methionine oxidation, methionine excision at the N terminus were selected as variable modifications. Results were filtered for a 1% false discovery rate (FDR) on spectrum, peptide, and protein levels. Relative protein abundances and pairwise comparisons were calculated using the MSstats package^46^ with default settings selected.

### CPRG lysis assay

Overnight cultures of *P. aeruginosa* and *V. cholerae*, were grown in LB at 37°C and 200 rpm without antibiotics. *A. baylyi* overnight cultures were grown at 30°C in similar conditions. Day cultures were launched in LB at the following dilutions: 1:100 for *P. aeruginosa*, 1:50 for *A. baylyi* and 1:400 for *V: cholerae*. After 2.5-3h, when all cultures reached an OD_600nm_ of 0.7-1.4, cultures were spun down at 12,000g for 2 min at room temperature. The cultures were concentrated to OD 1 and mixed in a 1:2 prey:attacker ratio. Three μL of the resulting suspensions were spotted in wells of a flat-bottom 96-well plate that had been extemporaneously prepared with 150 μL LB with 1% agar containing 0.1 mM IPTG (isopropyl-β-D-thiogalactoside) and 20 μg/mL CPRG (red-β-D-galactopyranoside) and pre-dried for 20 min at room temperature next to the Bunsen burner. The plates were incubated in an Epoch2 plate reader (BioTek) at 30°C, and the OD_575nm_ was measured every 10 min. The curves were plotted with Prism.

### Flow cytometry

Bacteria with different fluorescently labelled T6SS sheaths (TssB1-mNG or TssB2-mCh2) were grown to mid-exponential phase at 37°C from an overnight culture in LB. Prior to measurement, the samples were diluted 100x in PBS and were analysed in a BD Fortessa flow cytometer, with 950V green and 950V red laser in low flow rate mode. Data were analysed using FlowJo v10, and the median fluorescence intensity (MFI) was plotted using Prism.

### Growth curves

Overnight cultures of *P. aeruginosa* were diluted to OD_600nm_ 1 in LB, and used to inoculate wells, in triplicate, of a flat-bottom 96-well plate containing LB or LB without NaCl, with or without antibiotics, at a starting OD_600nm_ of 0.05. They were incubated at 37°C in an Epoch2 plate reader (BioTek) with orbital shaking at 240cpm and the OD_600nm_ was measured every 10 min. If antibiotic treatment was performed, antibiotics were added after 1.5h of incubation at 20 µM. The resulting curves were plotted in Prism.

### Fluorescence microscopy

Overnight cultures of *P. aeruginosa* were diluted 100-fold in LB and incubated for around 3h at 37°C and 200 rpm. They were then pelleted on a tabletop centrifuge at 12.000g for 2 min at room temperature and resuspended to OD 10 with LB containing 4 μg/mL FM4-64 (T13320, ThermoFisher Scientific) and incubated for 10 min at room temperature in the dark. One μL of the cell suspensions were then placed on agar pads (1/3 LB, 2/3 PBS with 1% agarose) and imaged on a Nikon Ti-E inverted motorized microscope with Perfect Focus System and Plan Apo 200x Oil Ph3 DM (NA 1.4) objective lens. Spectra X light engine (Lumencor) and ET-mCherry (Chroma 49008) filter sets were used to excite and filter fluorescence and a sCMOS camera pco.edge 4.2 (PCO, Germany, pixel size 65nm) and VisiView software (Visitron Systems, Germany) were used. Imaging was carried out at 30 °C by an Okolab T-unit (Okolab).

### Whole genome sequencing

Genomic DNA was extracted from overnight bacterial cultures in 1.5mL LB following the instructions of the GenElute Bacterial Genomic DNA kit (NA2110, Sigma). DNA concentration was measured using the Qubit dsDNA Quantitation Broad Range Assay kit (Q32850, ThermoFisher Scientific) and barcoding was performed according to the manufacturer’s instructions using the Rapid Barcoding Kit 24 V14 (SQK-RBK114.24, Oxford Nanopore Technologies). Samples were sequenced on a MinION flow cell (R10.4.1, FLO-MIN114, Oxford Nanopore Technologies). Basecalling was performed using Dorado software from Nanopore and the genomes were compared using the Snakemake evo-genome-analysis pipeline that is publicly available at https://github.com/mmolari/evo-genome-analysis/. This pipeline uses the raw reads from each sample and maps it to the reference using Minimap2^47^, extracting genomic changes including single nucleotide polymorphisms, gaps, insertions, clips and rearrangements.

### Statistics

When only two means were compared, we performed unpaired t-tests. When multiple means were compared, we used rdinary one-way analysis of variance (ANOVA) with Dunnett’s multiple comparison test, using GraphPad Prism version 9.3.1. Unless otherwise stated, data are represented as mean ± standard deviation (SD). *p-value<0.05, **p-value<0.005, ***p-value<0.0005, ****p-value<0.00005, ns – p-value>0.05.

## Acknowledgements

This work was supported by the National Center of Competence in Research AntiResist funded by the Swiss National Science Foundation (51NF40_180541). We thank Mattia Zampieri for providing the antibiotic collection used in this study.

## Data availability

All mass spectrometry raw data files associated with this manuscript are accessible at MassIVE (https://massive.ucsd.edu) under accession number MSV000096798. Genome sequencing data for this study have been deposited in the European Nucleotide Archive (ENA) at EMBL-EBI under accession number PRJEB84063 (https://www.ebi.ac.uk/ena/browser/view/PRJEB84063).

## Supplementary Figure legends

**Supplementary Figure 1.**
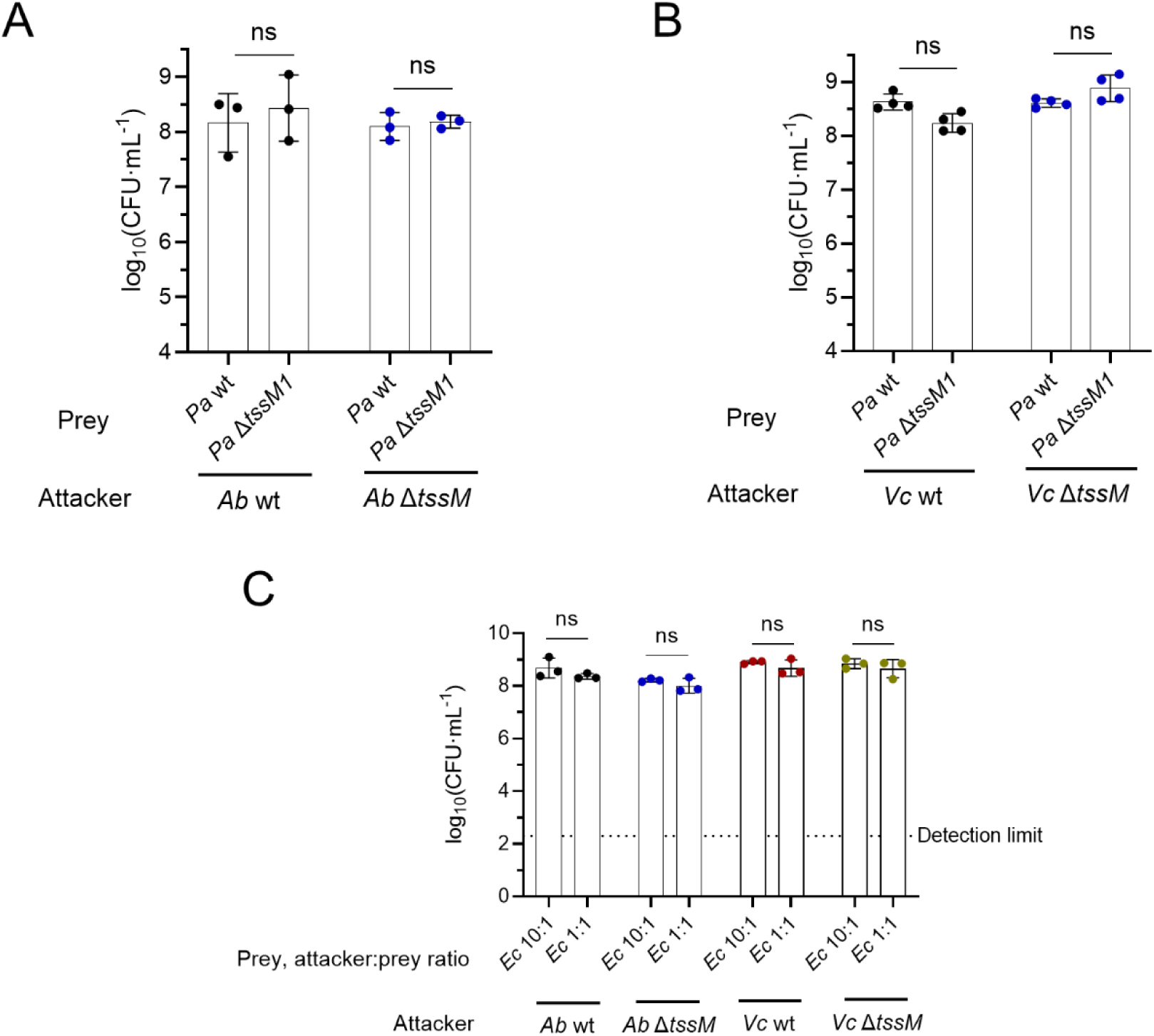
CFU counts showing the survival of the attacker when competing against *A. baylyi* (*Ab*) (A) and *V. cholerae* (*Vc*) (B) strains that have or not an active T6SS, in a 10:1 attacker:prey ratio where *P. aeruginosa* is the prey. (C) CFU counts showing the survival of *A. baylyi* and *V. cholerae* when competed against *E. coli* (*Ec*) in 10:1 and 1:1 attacker:prey ratios where *E. coli* is the prey.

**Supplementary Figure 2.**
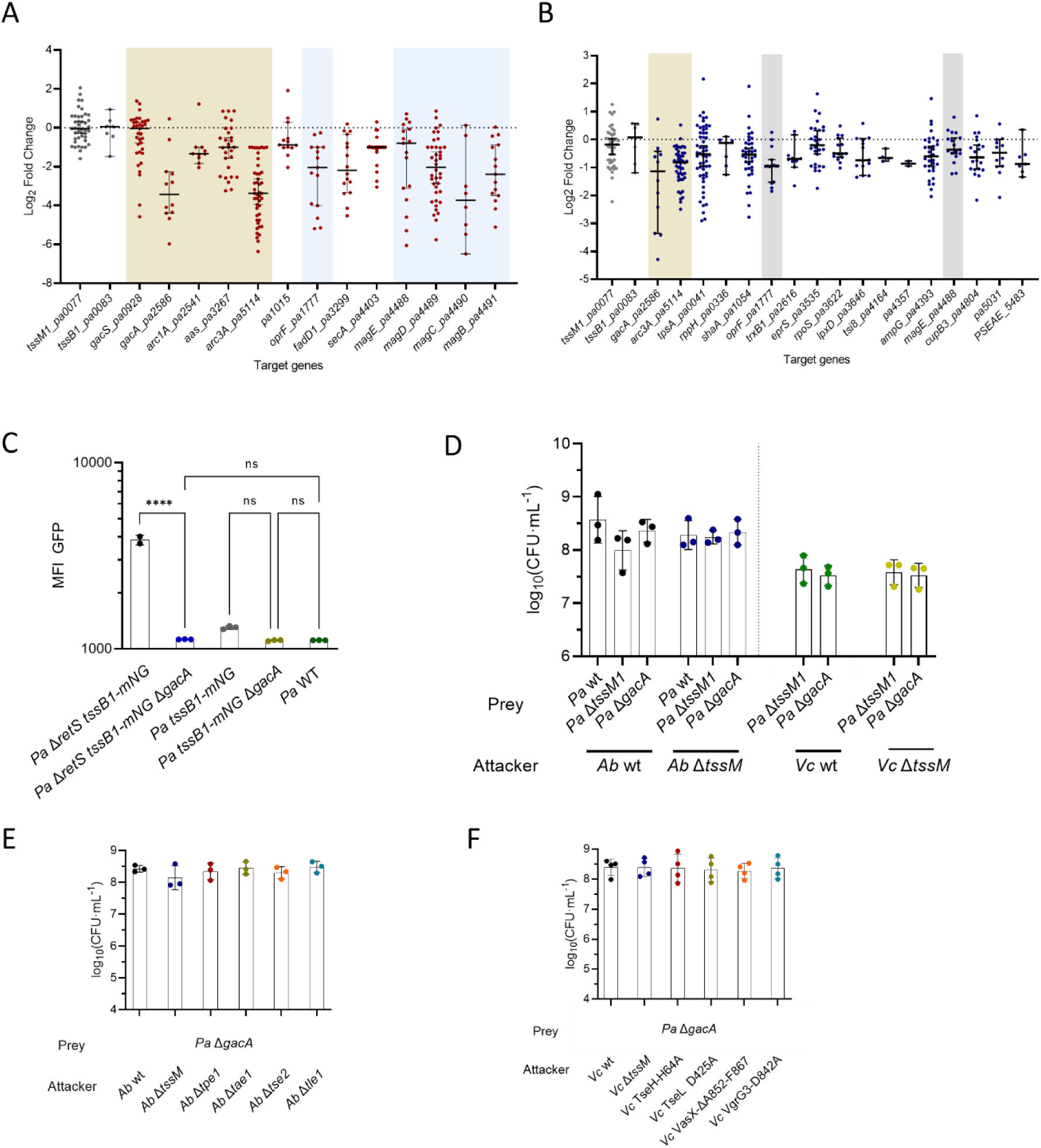
Log fold change of each sgRNA targeting the statistically significant hits of the CRISPRi screening where *P. aeruginosa* was competed against *V. cholerae* (A) or *A. baylyi* (B). (C) Median Fluorescence Intensity (MFI) of Green Fluorescent Protein (GFP) of wild type, Δ*retS* and Δ*gacA* strains where the sheath protein TssB1 of the H1-T6SS is labelled with mNeongreen where indicated. (D) CFU counts showing the survival of *A. baylyi* and *V. cholerae* when competed against *P. aeruginosa* strains in a 10:1 attacker:prey ratio where *P. aeruginosa* is the prey. (E) CFU counts of *A. baylyi* strains lacking different T6SS when competed against *P. aeruginosa* Δ*gacA* in a 10:1 ratio. (F) CFU counts of *V. cholerae* strains carrying different T6SS inactivated effectors when competed against *P. aeruginosa* Δ*gacA* in a 10:1 ratio.

**Supplementary Figure 3.**
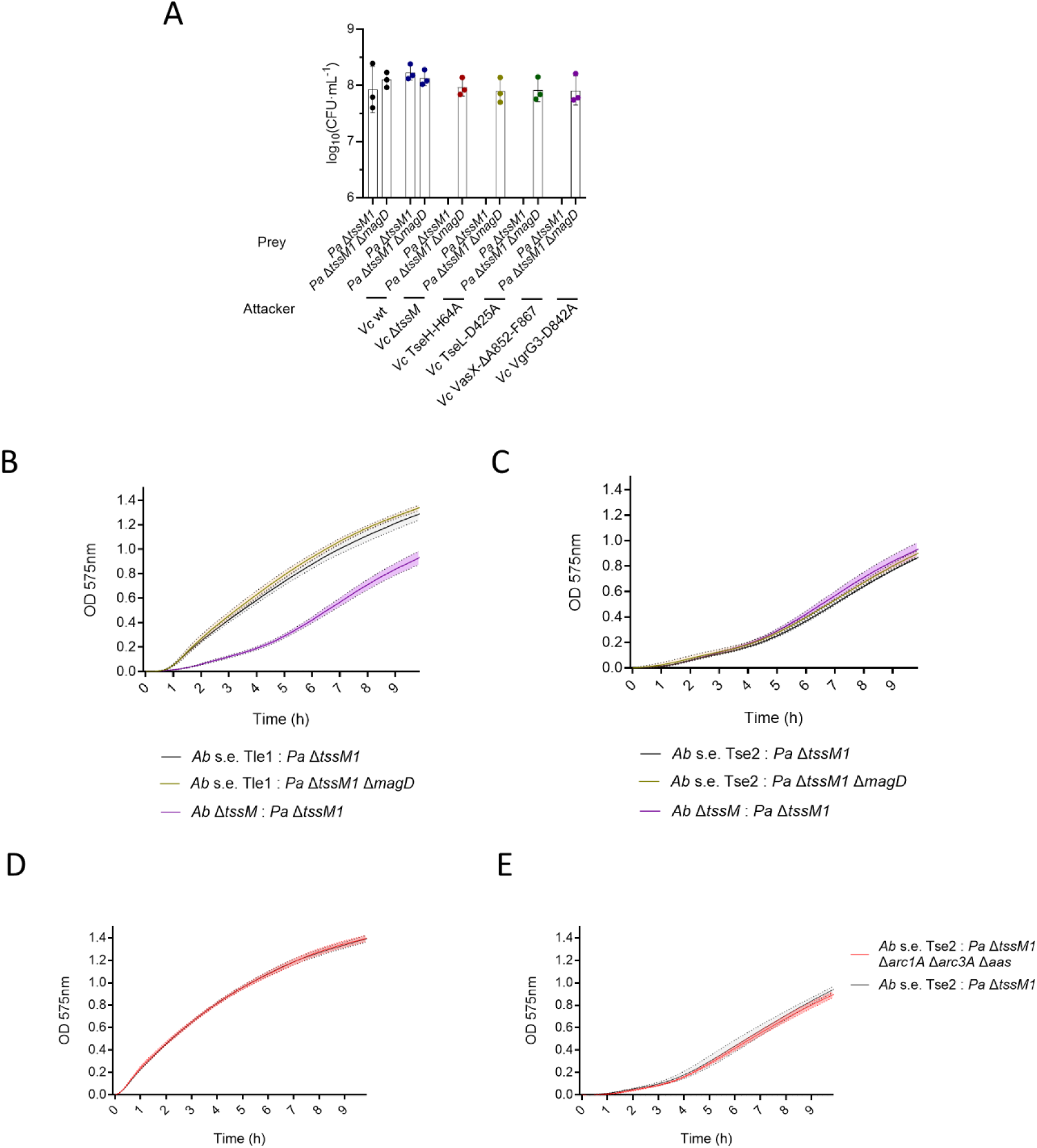
(A) CFU counts of *V. cholerae* strains carrying different T6SS inactivated effectors when competed against *P. aeruginosa* with or without *magD* in a 10:1 ratio. (B) CPRG lysis assay of *P. aeruginosa* with or without *magD*, competed against *A. baylyi* carrying only its lipase effector, Tle1. (C) CPRG lysis assay of *P. aeruginosa* with or without *magD*, competed against *A. baylyi* carrying only its Tse2 effector. (D) CPRG lysis assay of *P. aeruginosa* with or without *arc1A*, *arc3A* and *aas*, competed against *A. baylyi* carrying only its lipase effector, Tle1. (E) CPRG lysis assay of *P. aeruginosa* with or without *arc1A*, *arc3A* and *aas*, competed against *A. baylyi* carrying only its Tse2 effector.

**Supplementary Figure 4.**
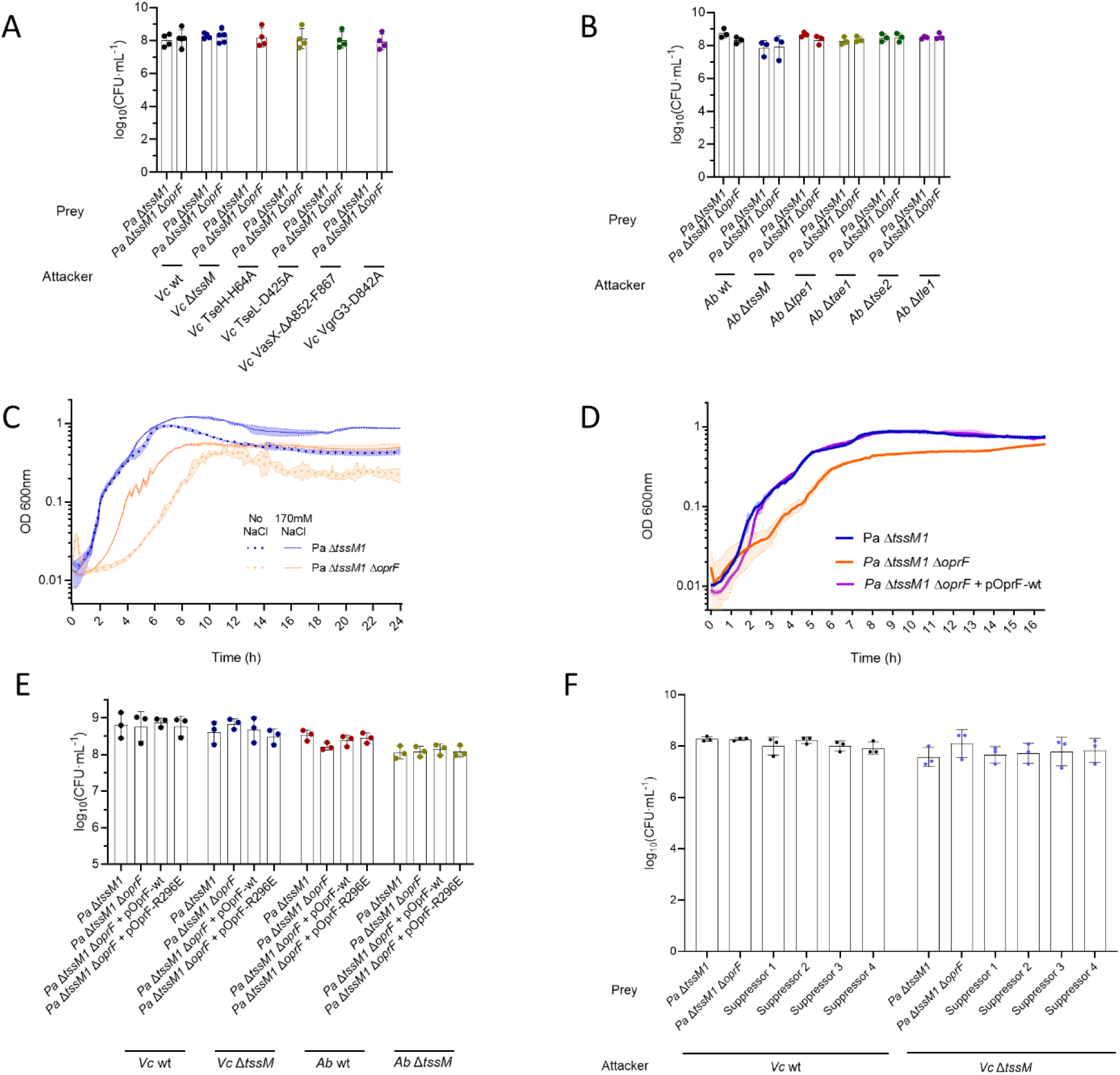
(A) CFU counts of *V. cholerae* strains carrying different T6SS inactivated effectors when competed against *P. aeruginosa* with or without *oprF* in a 10:1 ratio. (B) CFU counts of *A. baylyi* strains lacking different T6SS when competed against *P. aeruginosa* with or without *oprF* in a 10:1 ratio. (C) Growth curve of *P. aeruginosa* Δ*tssM1* and Δ*tssM1* Δ*oprF* strains in LB and LB without salt. (D) Growth curve of *P. aeruginosa* Δ*tssM1* Δ*oprF* strains complemented *in trans* with wild-type *oprF* in LB without salt. (E) CFU counts of *V. cholerae* and *A. baylyi* strains with or without an active T6SS when competed against *P. aeruginosa* with different *oprF* variants in a 10:1 ratio. (F) CFU counts of *V. cholerae* when competed against *P. aeruginosa* Δ*tssM1* Δ*oprF* suppressors in a 10:1 ratio.

**Supplementary Figure 5.**
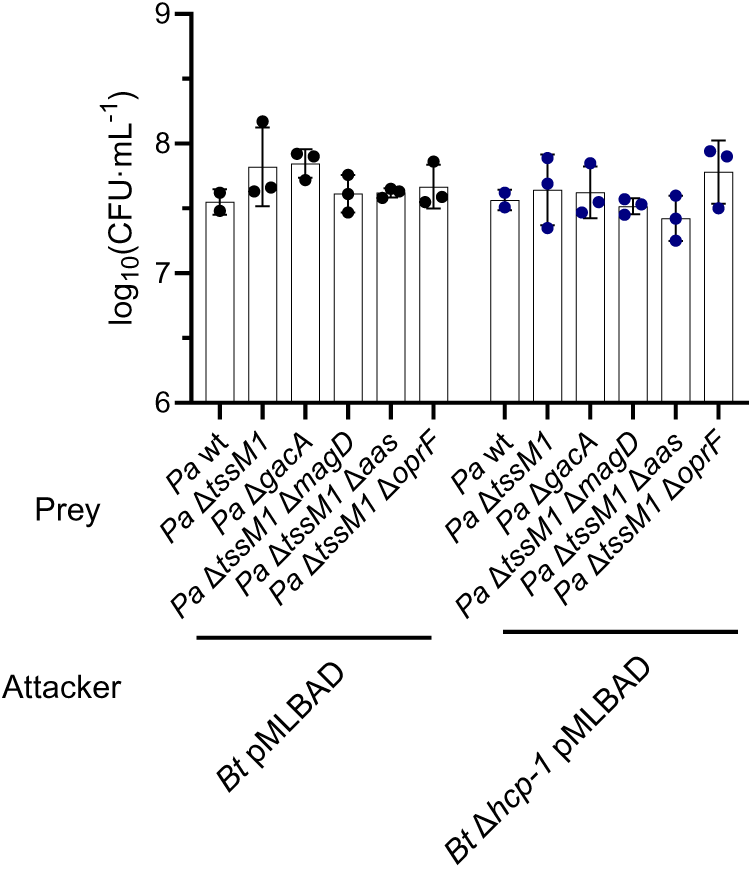
CFU counts of *B. thailandensis* with active or inactive T6SS-1 when competed against *P. aeruginosa* strains lacking different T6SS resistance mechanisms in a 10:1 ratio.

**Supplementary Figure 6.**
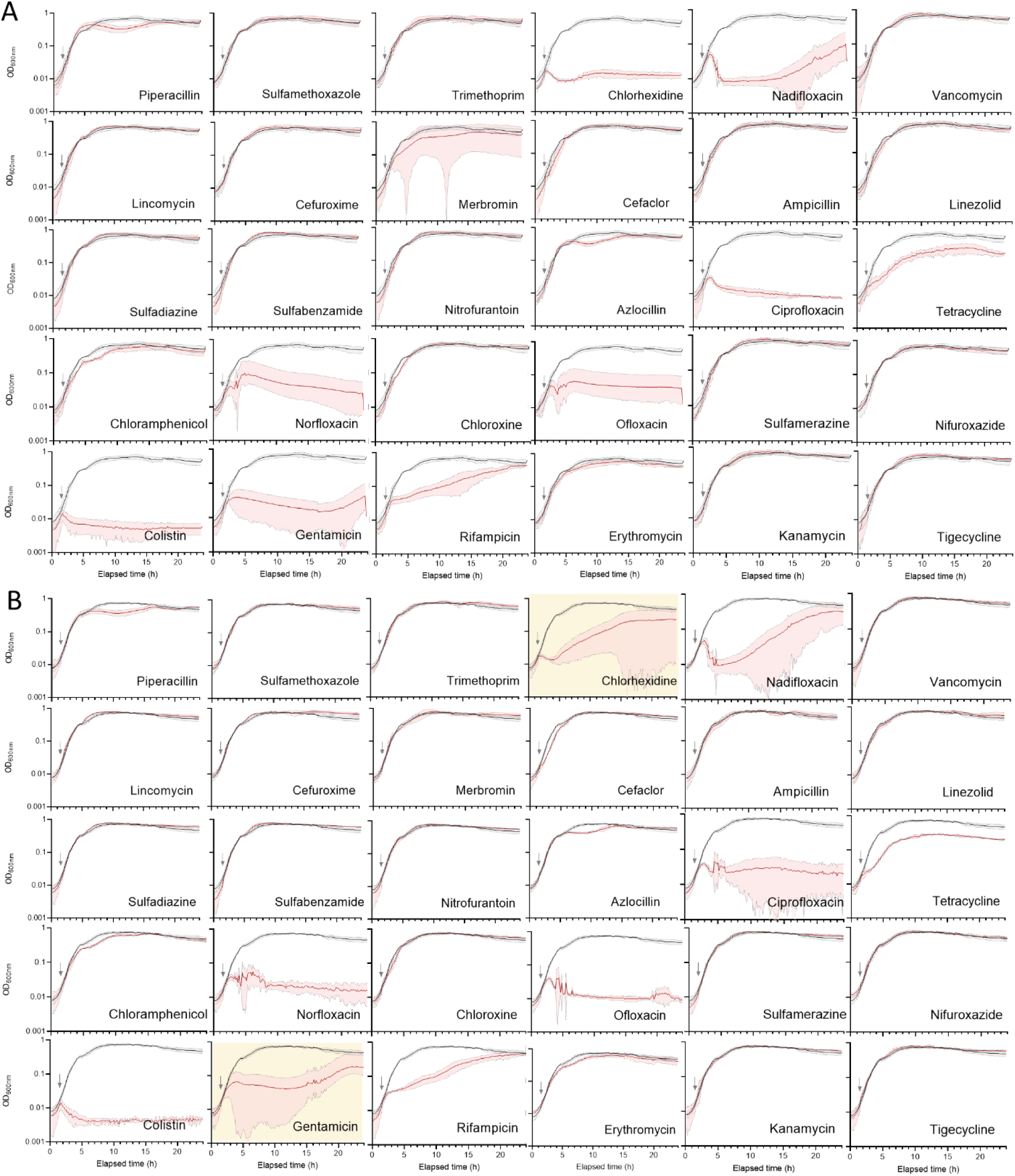
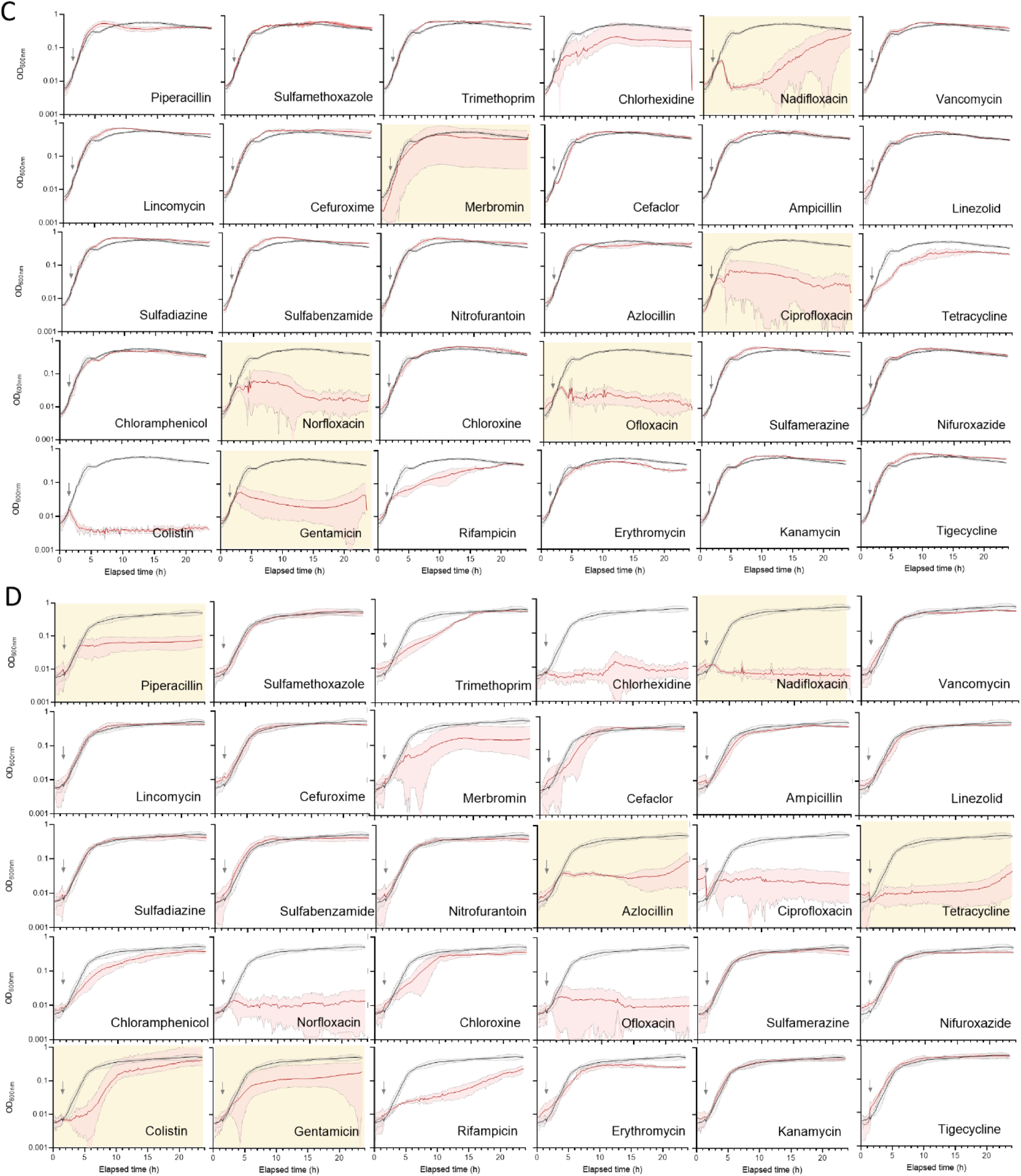
Growth curves of *P. aeruginosa* strains treated with a DMSO control (black curves) or different antibiotics (red curves) at a concentration of 20 μM. The grey arrow indicates the point at which antibiotics were added. (A) Curves for the wild-type parental strain. (B) Curves for the Δ*magD* strain. (C) Curves for the Δ*tssM1* Δ*arc1A* Δ*arc3A* Δ*aas* strain. (D) Curves for the Δ*oprF* strain. The treatments that show different behaviours compared to the parental strain are highlighted with a yellow background.

