## Supplementary Tables S2-S4 for "Mechanisms of *Pseudomonas aeruginosa* resistance to Type VI Secretion System attacks"

**Supplementary Table S2. Strains used in this study.**

| Strain number | Strain genotype | Reference |
| --- | --- | --- |
| *E. coli* | | |
| sATA016 | *E. coli* DH5α λpir | ^1^ |
| sATA380 | *E. coli* SM10 λpir | ^2^ |
| B252 | *E. coli* MG1665, GmR, *lacZ+* | ^3^ |
| *V. cholerae* 2740-80 Sm^R^, *lacZ*- | | |
| sATA075 | WT | ^4^ |
| sATA087 | Δ*tssM* | This study |
| sAP073 | *tseH-H64A* | This study |
| sAP077 | *vasX-ΔA852-F867* | This study |
| sAP078.2 | *vgrG3-D842A* | This study |
| sAP087 | *tseL-D425A* | ^5^ |
| *A. baylyi* ADP1 *rpsL-K88R*, *lacZ-* | | |
| sATA071 | WT | ^3^ |
| sATA694 | *vipA-mCh2* Δ*tssM* | This study |
| PR177 | *vipA-sfGFP clpV-mCh2* Δ*tpe1* | ^6^ |
| PR181 | *vipA-sfGFP clpV-mCh2* Δ*tae1* | ^6^ |
| DH055 | Δ*tse2* | ^6^ |
| DH056 | Δ*tle1* | ^6^ |
| sATA211 | *clpV-mCh2* Δ*tpe1* Δ*tae1* Δ*tse2* (single effector Tle1) | ^6^ |
| sATA212 | *clpV-mCh2* Δ*tpe1* Δ*tae1* Δ*tle1* (single effector Tse2) | ^6^ |
| sATA213 | *clpV-mCh2* Δ*tpe1* Δ*tse2* Δ*tle1* (single effector Tae1) | ^6^ |
| *P. aeruginosa* PAO1 | | |
| sATA001 | WT | ^7^ |
| sATA112 | Δ*tssM1* | This study |
| sATA250 | Δ*gacA* | This study |
| sATA681 | Δ*tssM1 attn7::dCas9* | This study |
|  | Δ*magD* | This study |
| sAP161 | Δ*tssM1* Δ*magD* | This study |
| sATA1117 | Δ*tssM1* Δ*magD* pME3856 (Ptac-LacZ) | This study |
| sAP218 | Δ*tssM1* Δ*aas* | This study |
| sATA1405 | Δ*tssM1* Δ*aas* pME3856 (Ptac-LacZ) | This study |
| sATA770.1 | Δ*tssM1* Δ*arc1A* | This study |
|  | Δ*tssM1* Δ*arc1A* pME3856 (Ptac-LacZ) | This study |
| sATA787 | Δ*tssM1* Δ*arc3A* | This study |
|  | Δ*tssM1* Δ*arc3A* pME3856 (Ptac-LacZ) | This study |
| sAP185 | Δ*oprF* | This study |
| sAP186 | Δ*tssM1* Δ*oprF* | This study |
| sATA1714 | Δ*tssM1* Δ*arc1A* Δ*arc3A* Δ*aas* | This study |
| sATA1718 | Δ*tssM1* Δ*arc1A* Δ*arc3A* Δ*aas* pME3856 (Ptac-LacZ) | This study |
| sATA1724 | Δ*tssM1* Δ*oprF* pPSV35-*oprF-R296E* | This study |
| sATA1733 | Δ*tssM1* Δ*oprF* pPSV35-*oprF* | This study |
| *B. thailandensis* E264 | | |
| MP002 | WT | ^8^ |
| sATA1660 | *tssB1-msfGFP* pMLBAD | This study |
| sATA1661 | *tssB1-msfGFP* Δ*hcp-1* pMLBAD | This study |

**Supplementary Table S3. Plasmids used in this study.**

| Plasmid | Antibiotic resistance | Reference |
| --- | --- | --- |
| pEXG2 | Gentamicin | ^9^ |
| pEXG2-Δ*tssM1* |  | ^5^ |
| pEXG2-Δ*gacA* |  | This study |
| pEXG2-Δ*magD* |  | This study |
| pEXG2-Δ*aas* |  | This study |
| pEXG2-Δ*arc1A* |  | This study |
| pEXG2-Δ*arc3A* |  | This study |
| pEXG2-Δ*oprF* |  | This study |
| pWM91 | Ampicillin | ^4,10^ |
| pWM91-Δ*tssM* |  | ^11^ |
| pWM91-Δ*tseH* |  | This study |
| pWM91-Δ*tseL* |  | ^5^ |
| pWM91-Δ*vasX* |  | This study |
| pWM91-Δ*vgrG3* |  | This study |
| pWM91-*tseH-H64A* |  | This study |
| pWM91-*tseL-D425A* |  | ^5^ |
| pWM91-*vasX-ΔA852-F867* |  | This study |
| pWM91-*vgrG3-D842A* |  | This study |
| pME3856 | Tetracycline | ^12^ |
| pPSV35 | Gentamicin | ^9^ |
| pPSV35-*oprF* |  | This study |
| pPSV35-*oprF-R296E* |  | This study |
| pMLBAD | Trimethoprim | ^13^ |
| pUC18T:TNT7:dCas9Spas | Gentamicin | (Kaczmarczyk *et al.*) |
| pTNS2 | Gentamicin |  |
| pFLP2 | Carbenicillin |  |

**Supplementary Table S4. Primers used in this study.**

| Primer | Sequence | Description |
| --- | --- | --- |
| oATA117 | GTACCGAATTCGAGCTCGAGCCCGGGGATCCTCTAGAggaggagcccgaggacgagg | fw primer to amplify 800bp from the region upstream from GacA (PA2586), with an overlap to the XbaI site from the pEXG2 plasmid |
| oATA118 | gaccaccagcaccttaatcacg | rv primer to amplify the region upstream from GacA (PA2586), including the first 7 aa |
| oATA119 | tgattaaggtgctggtggtcggcatggtcgatgccgccagc | fw primer to amplify the region downstream from GacA (PA2586), with an overlap to oATA118 |
| oATA120 | TATACGAGCCGGAAGCATAAATGTAAAGCAAGCTTcgccgcgtacgctgatcagg | rv primer to amplify 800bp from the region downstream from GacA (PA2586), with an overlap to the HindIII site from the pEXG2 plasmid |
| oATA133 | gtgcgacgttatgtctggcc | fw primer to check the deletion of gacA (PA2586) |
| oATA134 | tcacgttgacgatcaactcc | rv primer to check the deletion of gacA (PA2586) |
| P02682 | dPA4489_1_HindIII.FOR | TCAGTAAAGCTTCCGGAAGCCAGGCCAACGC |
| P02683 | dPA4489_1.REV | CAGCGCCGGGCTACGGCTGAAACGGC |
| P02684 | PA4489_2.FOR | AGCCGTAGCCCGGCGCTGGCCAAGGT |
| P02685 | dPA4489_2_Xba1.REV | TCAGTATCTAGAGCACCTCGGCGTAGCGGTC |
| P02686 | dPA4489_Det.FOR | CCTGGACACCCTCGACAAA |
| P02687 | dPA4489_Det.REV | ACCCGTTGCAGGAAGAGG |
| oAP109 | TATACGAGCCGGAAGCATAAATGTAAAGCAAGCTTccaccaagcaccagggcatc | 700bp upstream of PA3267, with HindIII top strand |
| oAP110 | gaattgcgaatgttgggtcat | first 7aa of PA3267 |
| oAP111 | acccaacattcgcaattccgcggcgaagctcgctga | last 7aa of PA3267 and complementary overlap with oAP110 |
| oAP112 | GTACCGAATTCGAGCTCGAGCCCGGGGATCCTCTAGActtcagcaagccgcttccc | ca 700bp downstream of PA3267 , with XbaI bottom strand |
| oAP113 | gccaagacctacgaactggg | Deletion verification of PA3267 (upstream) |
| oAP114 | ccggtgatcgaatggggacg | Deletion verification of PA3267 (downstream) |
| oATA370 | TATACGAGCCGGAAGCATAAATGTAAAGCAAGCTTgatggacatcgatgtcgagg | fw primer to amplify 800 bp upstream from PA2541, arc1A, to make a deletion strain, with an overlap to clone by Gibson assembly into the HindIII restriction site of pEXG2 plasmid |
| oATA371 | ttgatagacggaaatcatgtcagc | rv primer to amplify the first 7 codons of PA2541, arc1A, to make a deletion strain |
| oATA372 | acatgatttccgtctatcaaggcagcgacggccaggcc | fw primer to amplify the last 7 codons of PA2541, arc1A, to make a deletion strain, with an overlap to primer oATA371 |
| oATA373 | GTACCGAATTCGAGCTCGAGCCCGGGGATCCTCTAGAgctcctgcggcgcacgatgg | rv primer to amplify 800 bp downstream from PA2541, arc1A, to make a deletion strain, with an overlap to clone by Gibson assembly into the XbaI restriction site of the pEXG2 plasmid |
| oATA374 | ctatgcgcaactggaggtgg | fw primer to assess the deletion of PA2541, arc1A |
| oATA375 | agcaggctcggtcgttcagg | rv primer to assess the deletion of PA2541, arc1A |
| oATA382 | TATACGAGCCGGAAGCATAAATGTAAAGCAAGCTTtatgtccagttgccgcatgc | fw primer to amplify 800 bp upstream from PA5114, arc3A, to make a deletion strain, with an overlap to clone by Gibson assembly into the HindIII restriction site of pEXG2 plasmid |
| oATA383 | cagcatgaaaatccattgcatgc | rv primer to amplify the first 7 codons of PA5114, arc3A, to make a deletion strain |
| oATA384 | tgcaatggattttcatgctgtcggaggccgagcagccatgac | fw primer to amplify the last 7 codons of PA5114, arc3A, to make a deletion strain, with an overlap to primer oATA383 |
| oATA385 | GTACCGAATTCGAGCTCGAGCCCGGGGATCCTCTAGAgctgcaacggcaacgccagc | rv primer to amplify 800 bp downstream from PA5114, arc3A, to make a deletion strain, with an overlap to clone by Gibson assembly into the XbaI restriction site of the pEXG2 plasmid |
| oATA386 | tgccggcaatctcgtccagc | fw primer to assess the deletion of PA5114, arc3A |
| oATA387 | ctcgttctgctgtacgttgc | rv primer to assess the deletion of PA5114, arc3A |
| oAP088 | GTACCGAATTCGAGCTCGAGCCCGGGGATCCTCTAGAatatgactggcctgatctca | ca 700bp upstream of oprF (PA1777), with XbaI bottom strand |
| oAP089 | gcctaaggtgttcttcagtttcat | first 8aa of oprF (PA1777) |
| oAP090 | aaactgaagaacaccttaggcgtagaagctgaagccaagtaa | last 7aa of oprF (PA1777) and complementary overlap with oAP089 |
| oAP091 | TATACGAGCCGGAAGCATAAATGTAAAGCAAGCTTtggtgatggtcgacgatctg | ca 700bp downstream of oprF (PA1777), with HindIII top strand |
| oAP092 | gacggaagagggagagcttg | deletion verification of oprF (upstream) |
| oAP093 | atggcagcgaatgcctgagg | deletion verification of oprF (downstream) |
| oATA881 | ATGACCATGATTACGAATTCAGGAGGAAACTAGTatgaaactgaagaacaccttaggc | fw primer to amplify PA1777_OprF with an overlap for insertion in the EcoRI site of pPSV35 and an RBS that also contains a stop codon for the lacZa gene that starts translation in the IPTG-inducible promoter |
| oATA885 | AGGTCGACTCTAGAGGATCCttacttggcttcagcttctacttcggcttc | rv primer to amplify PA1777_OprF with an overlap for insertion in the BamHI site of pPSV35 and a STOP codon |
| oATA779 | acgtcacgaacggcgttggcctcacgctcggacagcttctggttg | rv primer to amplify the region downstream from R296 in oprF, PA1777; to introduce mutation R296E |
| oATA851 | cgtgaggccaacgccgttcg | fw primer to amplify the region downstream from R296 in oprF, PA1777 |
| oAP001 | tcactcagataccgttaattttggagttgatgatacagtgtcagtcattag | 5' end of TseH (VCA0285) with complementary overlap of oAP002 |
| oAP002 | aacaaattaacggtatctgagtgatatg | 3' end of TseH (VCA0285) |
| oAP003 | CGGCCGCTCTAGAACTAGTgtcgctttcgcaagacctgggccg | rv primer for dTseH (VCA0285) with SpeI bottom strand |
| oAP004 | CGAATTCCTGCAGCCCGGgtcacaaaatcaattagctggtgaacgg | fw primer for dTseH (VCA0285) with XmaI top strand |
| oAP011 | gctggagttattttgattaatg | TseH (VCA0285) point mutation H64A primer 1 (top w/o mutation) |
| oAP012 | cattaatcaaaataactccagccggaccaaggtagggggc | TseH (VCA0285) point mutation H64P primer 2 (bottom w/ mutation) |
| oAP023 | catgtgatgcattatgttcaactc | top strand 100 bp upstream of TseH (VCA0285)/TsiH insertion (for Vc integration verification) |
| oAP024 | catgtggggtacgtgtggacc | bottom strand 100 bp downstream of TseH (VCA0285)/TsiH insertion (for Vc integration verification) |
| oAP015 | acactcttaggtagtggaattg | VasX (VCA0020) deletion A852-F867 |
| oAP016 | caattccactacctaagagtgtttttaacaaataaccgaagcg | VasX (VCA0020) deletion A852-F868 W/ overlap oAP015 |
| oAP017 | CGGCCGCTCTAGAACTAGgcctttgtacctttttccgc | bottom 700bp upstream VasX (VCA0020) with SpeI bottom strand |
| oAP018 | CGAATTCCTGCAGCCCGGctattcctcttttaattcttg | top 700bp downstream VasX (VCA0020) with XmaI top strand |
| oAP025 | catcctcggcttttacgtcagtc | top strand 100 bp downstream of VasX (VCA0020)/TsiV-2 insertion (for Vc integration verification) |
| P02742 | GCTGTACAAAgaggccaagggaaattaatgc | Bottom strand upstream of VasX (VCA0020)/TsiV-2 insertion (for Vc integration verification) |
| oAP013 | tatggtggggcttcatatggttg | VgrG3 (VCA0123) point mutation D842A primer 1 (top w/o mutation) |
| oAP014 | catatgaagccccaccataaactccatttcctgttgatataacc | VgrG3 (VCA0123) point mutation D842V primer 2 (bottom w/ mutation) |
| oAP019 | CGAATTCCTGCAGCCCGGcaacatgatctgatcccgctgtatc | top 700 bp upstream VgrG3 (VCA0123) with XmaI top strand |
| oAP020 | CGGCCGCTCTAGAACTAGTgcgaagtgggcagcaaagcttg | bottom 700 bp downstream VgrG3 (VCA0123) with SpeI bottom strand |
| oATA402 | tcgtcggcagcgtcagatgtgtataagagacagtcagtgatagagatactgagcac | Primer 16587_sgRNA_par0_F from (Kaczmarczyk *et al.*) |
| oATA403 | gtctcgtgggctcggagatgtgtataagagacagtctttgtaggctctttcgagtac | Primer 16591_sgRNA_par0_R from (Kaczmarczyk *et al.*) |
| oATA404 | tcgtcggcagcgtcagatgtgtataagagacagctcagtgatagagatactgagcac | Primer 16588_sgRNA_par1_F from (Kaczmarczyk *et al.*) |
| oATA405 | gtctcgtgggctcggagatgtgtataagagacagctctttgtaggctctttcgagtac | Primer 16592_sgRNA_par1_R from (Kaczmarczyk *et al.*) |
| oATA406 | tcgtcggcagcgtcagatgtgtataagagacagactcagtgatagagatactgagcac | Primer 16589_sgRNA_par2_F from (Kaczmarczyk *et al.*) |
| oATA407 | gtctcgtgggctcggagatgtgtataagagacagactctttgtaggctctttcgagtac | Primer 16593_sgRNA_par2_R from (Kaczmarczyk *et al.*) |
| oATA408 | tcgtcggcagcgtcagatgtgtataagagacaggactcagtgatagagatactgagcac | Primer 16590_sgRNA_par3_F from (Kaczmarczyk *et al.*) |
| oATA409 | gtctcgtgggctcggagatgtgtataagagacaggactctttgtaggctctttcgagtac | Primer 16594_sgRNA_par3_R from (Kaczmarczyk *et al.*) |
